# G6b-B antibody-based *cis*-acting platelet receptor inhibitors (CAPRIs) as a new family of anti-thrombotic therapeutics

**DOI:** 10.1101/2024.05.10.593500

**Authors:** Alexandra Mazharian, Ophélie Bertin, Amrita Sarkar, Johanna Augros, Alicia Bornert, Cécile Loubière, Friederike Jönsson, Juli Warwicker, W. Abbott Mark, Omer Dushek, Caroline Vayne, Lubica Rauova, Klaus Fütterer, Jérôme Rollin, Mortimer Poncz, Yotis A. Senis

## Abstract

**Key Points:**

- Bispecific *cis-*acting platelet receptor inhibitors (CAPRIs) mediate hetero-clustering of the ITIM receptor G6B with ITAM receptors.
- G6B-GPVI and G6B-CD32A CAPRIs specifically inhibit collagen- and immune complex-induced thrombus formation, respectively.

Platelets are highly reactive fragments of megakaryocytes that play a fundamental role in thrombosis and hemostasis. Predictably, all conventional anti-platelet therapies elicit bleeding, raising the question whether the thrombotic activity of platelets can be targeted separately. In this study, we describe a novel approach of inhibiting platelet activation through the use of bispecific single-chain variable fragments (bi-scFvs), termed *cis-*acting platelet receptor inhibitors (CAPRIs) that harness the immunoreceptor tyrosine-based inhibition motif (ITIM)-containing co-inhibitory receptor G6b-B (G6B) to suppress immunoreceptor tyrosine-based (ITAM)-containing receptor-mediated platelet activation. CAPRI-mediated hetero-clustering of G6B with either the ITAM-containing GPVI-FcR γ-chain complex or FcγRIIA (CD32A) inhibited collagen- or immune complex-induced platelet aggregation. G6B-GPVI CAPRIs strongly and specifically inhibited thrombus formation on collagen under arterial shear, whereas G6B-CD32A CAPRI strongly and specifically inhibited thrombus formation to heparin-induced thrombocytopenia, vaccine-induced thrombotic thrombocytopenia and antiphospholipid syndrome complexes on Von Willebrand Factor-coated surfaces and photochemical-injured endothelial cells under arterial shear. Our findings provide proof-of-concept that CAPRIs are highly effective at inhibiting ITAM receptor-mediated platelet activation, laying the foundation for a novel family of anti-thrombotic therapeutics with potentially improved efficacy and fewer bleeding outcomes compared with current anti-platelet therapies.

## Introduction

Platelets are highly reactive fragments of megakaryocytes (MKs) that play a fundamental role in thrombosis and hemostasis. Rapid platelet adhesion and activation at sites of vascular injury is essential for the formation of the primary hemostatic plug and localized thrombin generation, which consolidates the thrombus through fibrin deposition. Primary and secondary waves of platelet activation, adhesion and aggregation are mediated by a diverse repertoire of tyrosine kinase-linked and G protein-couple receptors, several of which are effective targets for inhibiting thrombosis, including the fibrinogen integrin αIIbβ3 (abciximab),(1) the ADP receptor P2Y_12_ (clopidogrel) and the thrombin protease activating receptor-1 (vorapaxar).(2, 3) The immunoreceptor tyrosine-based activation motif (ITAM)-containing collagen receptor complex GPVI-FcR γ-chain (glenzocimab), and von Willebrand factor (VWF) receptor complex glycoprotein (GP)Ib-IX-V (anfibatide) are emerging as promising antithrombotic drug targets.(4, 5) However, all conventional anti-platelet therapies targeting activation receptors and associated signaling pathways have bleeding side-effects, raising the need for new and improved ways of targeting platelets.

In this study, we describe the design and characterization of bispecific single-chain variable fragments (bi-scFvs), termed *cis-*acting platelet receptor inhibitors (CAPRIs) that harness the immunoreceptor tyrosine-based inhibition motif (ITIM)-containing MK and platelet co-inhibitory receptor G6B (MPIG6B, G6b-B, G6B) to inhibit signaling from the ITAM receptors, GPVI-FcR γ-chain receptor complex and immunoglobulin (Ig) FcγRIIA (CD32A) receptor. G6B is highly and specifically expressed on the MK/platelet lineage and plays a vital role in regulating platelet production and function.(6–8) It binds and is regulated by the heparan sulfate (HS) side-chains of perlecan in the vessel wall and structurally-related heparin.(9) Phosphorylation of tyrosine residues in the G6B ITIM and immunoreceptor tyrosine-based switch motif (ITSM) by Src family kinases (SFKs) facilitates binding and activation of the tandem Src homology 2 (SH2) domain-containing protein-tyrosine phosphatases 1 and 2 (Shp1 and Shp2) that are essential for the inhibitory activity of G6B.(10) In addition, phospho (p)-ITIM/ITSM-associated Shp1 and Shp2 must be within the molecular reach of components of the ITAM receptor proximal signaling complex, comprised of SFKs and p-ITAM-associated spleen tyrosine kinase (Syk) that can be dephosphorylated and inactivated.(11)

We hypothesized that physiological ligand-dependent hetero-clustering of ITIM-ITAM receptors can be induced using synthetic CAPRIs that will inhibit platelet activation. Findings from this study demonstrate this to be the case, and that bispecific G6B-GPVI and G6B-CD32A CAPRIs inhibit GPVI- or CD32A-induced platelet function, respectively. G6B-GPVI CAPRIs strongly and specifically inhibited thrombus formation on collagen under arterial shear, whereas the G6B-CD32A CAPRI strongly and specifically inhibited thrombus formation in three prothrombotic disorders involving platelet Fc-receptor activation: heparin-induced thrombocytopenia (HIT),(12) vaccine-induced thrombotic thrombocytopenia (VITT)(13) and antiphospholipid syndrome (APS) complexes on VWF-coated surfaces and photochemical-injured endothelial cells in microfluidic systems.(14, 15) G6B-GPVI CAPRIs had little or no effect at inhibiting thrombus formation involving HIT, VITT and APS immune complexes under flow, and conversely, G6B-CD32A CAPRI did not inhibit thrombus formation on collagen under flow, providing proof-of-concept of the specificity and effectiveness of CAPRIs at inhibiting ITAM receptor-mediated platelet activation and thrombosis.

## Materials and Methods

### Reagents

Anti-human platelet factor 4 (PF4) monoclonal antibodies (mAbs) KKO, 5B9 and 1E12 were generated as previously described.(16–18) Anonymized serum samples were obtained from patients diagnosed with HIT, VITT or APS. IgGs were isolated using protein G agarose (Thermo Fisher). Other reagents and antibodies are described in detail in Supplemental Materials.

### Production of bi-scFv CAPRIs

Bi- and mono-specific scFvs were generated as described in detail in Supplemental Materials.

### Structural modeling

Structural models of the receptor bridging complexes were constructed using a combination of AlphaFold2-generated structural models, the crystal structure of the G6b-B ectodomain bound to the 17.4 antibody (9), and manual modelling in ChimeraX (19). AlphaFold2 (20) was used to generate structural models of full-length receptors platelet glycoprotein VI (Uniprot Q9HCN6), G6b-B (Uniprot O95866) and FcgRIIA (Uniprot P31995), using the colab web interface (https://colab.research.google.com/github/sokrypton/ColabFold/blob/main/AlphaFold2.ipynb).

Similarly, scFv-ectodomain complex structures were constructed using co-folding sequences of receptor ectodomains and cognate scFv fragments in AlphaFold2, except for the G6b-B:17.4 complex. The resulting complexes were superimposed over the AF2-generated model of the bi-specific scFv fragments, followed by manual rotation of the TM helices vs. the globular ectodomains to achieve an approximate alignment of the TM helices.

### Platelet preparation

Blood was collected from healthy donors into anticoagulant citrate-dextrose solution (ACD), as previously described.(21) Washed human platelets were prepared using a standard protocol (21) Platelet counts were normalized and used for aggregation (3 x 10^8^/ml) or biochemical analysis (5 x 10^8^/ml).

### Platelet functional assays and biochemistry

Platelet aggregation, stimulation with HIT and VITT immune complexes, and for biochemical analysis are described in the Supplemental Methods.

### Immunoblotting

Platelet whole cell lysates (WCLs) were prepared and analyzed by automated capillary-based immunoassay (ProteinSimple Jess), prepared according to manufacturer’s instructions, as previously described (22) and in the Supplemental Methods. The High Dynamic Range (HDR) profile was used for chemiluminescent and fluorescent multiplexing signal detection. Optimized antibody dilutions and sample concentrations used are provided in Supplemental Table 2.

### Collagen microfluidic thrombosis studies

Platelet adhesion and thrombus formation was measured using a microfluidic system, as previously described.(23) Briefly, polydimethylsiloxane (PDMS) flow chambers channels (0.1 × 1mm) were coated with collagen (200 µg/ml) for 1 hour at room temperature and blocked with phosphate-buffered saline (PBS)-containing human serum albumin (10 mg/ml) for 30 minutes at room temperature. Whole blood was collected from healthy donors into 525 U/ml hirudin and treated with 0.2 µM of the various (bi-)scFvs for 15 minutes at 37°C prior to being perfused through the collagen-coated chamber with a syringe pump (Harvard Apparatus, Holliston, MA, USA) at 1,500 s^-1^ for 5 minutes at 37°C. Platelets adhesion and aggregation were visualized in real-time by differential interference contrast (DIC) microscopy (Leica DMI4000B). Platelets were also stained with 3,3′-dihexyloxacarbocyanine iodide (DiOC_6_) and visualized by confocal microscopy. Surface coverage by platelet aggregates (>100 µm²) and thrombus volume were quantified using ImageJ software. All experiments were performed blind.

### HIT and VITT microfluidic thrombosis studies on VWF

Microfluidic thrombosis models were performed as previously described.(17) Briefly, whole blood collected on 0.129 M sodium citrate from healthy donors, was pre-incubated with 0.1 µM of the various (bi-)scFvs for 10 minutes, prior to incubation with 1E12 at 10 µg/mL for 6 minutes. Blood samples were then recalcified to 2.7 mM CaCl_2_ and perfused at a shear rate of 20 µL/min (500 s^-1^; 20 dyn/cm²) in microfluidic channels (Vena8 Fluoro+, Cellix), pre-coated overnight at 4°C with 120 µg/mL purified human VWF (LFB). For more details, see the Supplementary Methods.

### Endothelial injury HIT, VITT and APS microfluidic studies

Microfluidic studies in 48-well BioFlux plates were performed with channels coated with near-confluent human umbilical vein endothelial cells (HUVECs, Lifeline Cell Technology), as previously described.(18) Citrated and calcein-labelled whole blood, supplemented with recombinant PF4 (25 μg/ml) and either 1 mg/ml of HIT, VITT or APS IgGs isolated from patient samples, in the presence or absence of 0.1 µM of the various (bi-)scFvs was perfused over the injured endothelium at 10 dyne/cm^2^. Platelet accumulation on the injured endothelium field was monitored. For more details, see the Supplementary Methods.

### Statistical analysis

Statistical analyses were performed with GraphPad Prism version 9 software. Statistical significance was analyzed by one- or two-way ANOVA followed by the appropriate *post hoc* test, or Kruskal-Wallis test for nonparametric data, a Bonferroni correction was applied for multiple comparisons, as indicated in figure legends. Differences were considered significant when the P values were <0.05.

### Institutional approval

Human sample studies were obtained after approval of the Children’s Hospital of Philadelphia’s institutional review board and were in accord with the Helsinki Principles. Legal and ethical authorization for the use of blood collected for research was obtained through a national convention between the French National Institute of Health and Medical Research (INSERM) and the French Blood Institute (EFS) (convention number I/DAJ/C2675) and conformed to the ethical standards of the Declaration of Helsinki.

## Results

### Development of bispecific ITIM-ITAM receptor CAPRIs

Inhibition of ITAM receptors by ITIM receptors is ligand- and proximity-dependent.(24) The primary aim of this study was to develop an antibody-based approach that mimics physiological ligand-mediated hetero-clustering of ITIM and ITAM receptors. G6B was the ITIM receptor of choice, because it is highly and specifically expressed in the MK/platelet lineage and it is an established inhibitor of platelet ITAM receptors.(6, 11) Bispecific G6B-GPVI and G6B-CD32A CAPRIs were engineered using well-characterized mAbs to the ectodomains of G6B, GPVI and CD32A. Variable sequences of anti-G6B and -GPVI mAbs 17.4 and 3J24 were used to generate the G6B-GPVI v1 CAPRI that binds outside the ligand binding sites of G6B and GPVI, respectively **(Figures 1A and 1B)**,(9, 25) whereas variable sequences of glenzocimab and anti-CD32A mAb IV.3 were used to generate the G6B-GPVI v2 and G6B-CD32A CAPRIs that bind within the ligand-binding sites of GPVI and CD32A, respectively.(5, 26) Therefore, the G6B-GPVI v1 CAPRI should inhibit GPVI signaling by hetero-clustering with G6B, whereas the G6B-GPVI v2 and G6B-CD32A CAPRIs should inhibit both ligand binding, as well as hetero-clustering with G6B **(Figure 1A)**. VH and VL sequences of antibodies were recombinantly connected using short G/S-rich linker sequences **(Figure 1B and Supplemental Figures 1A-1F)**, to provide the necessary flexibility and length to mediate G6B-GPVI and G6B-CD32A co-clustering, and be non-immunogenic.

**Figure 1.**
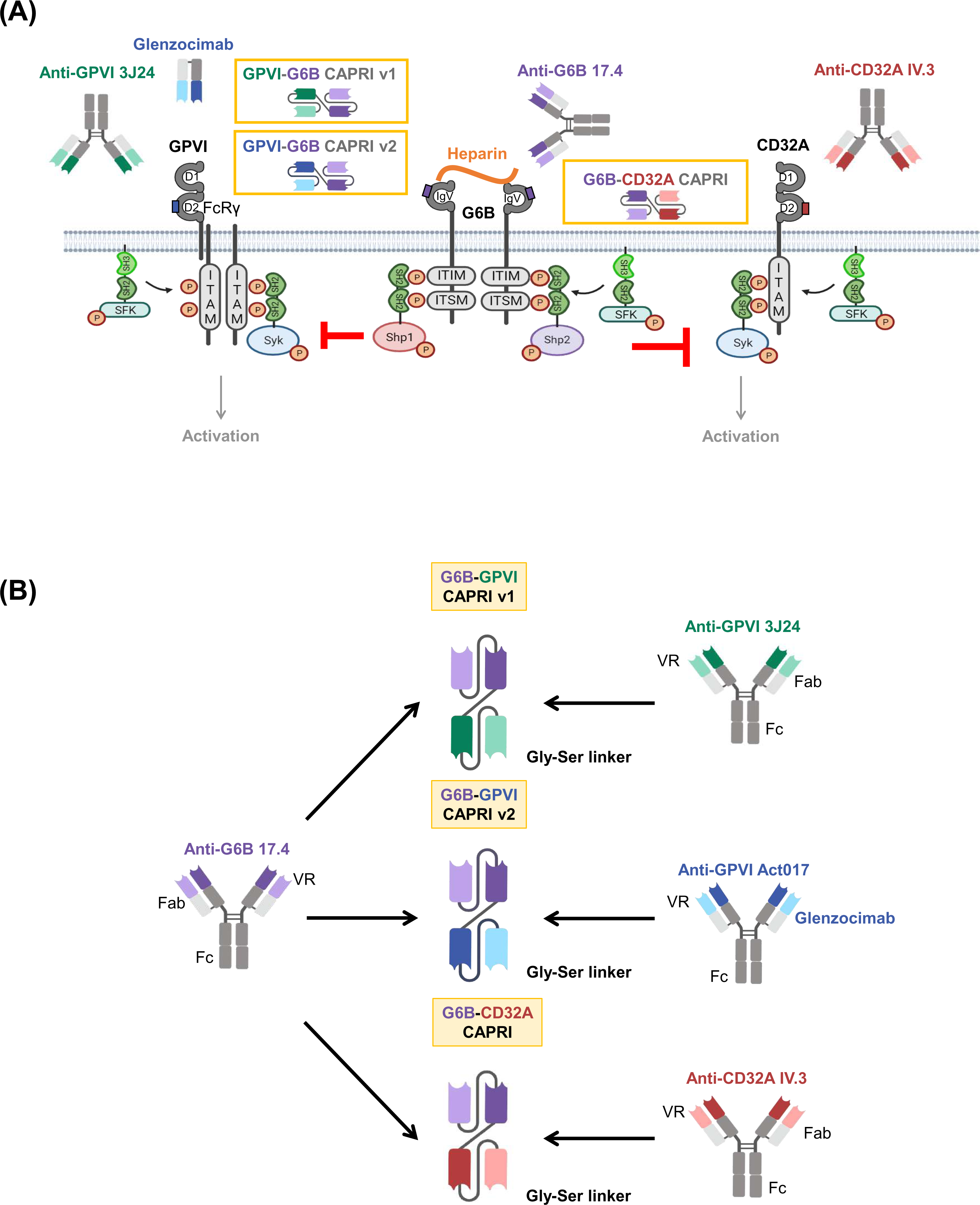
CAPRI-mediated ITIM-ITAM-containing receptor co-clustering. **(A)** The co-inhibitory receptor G6B consists of a single IgV ectodomain, and ITIM and ITSM in its cytoplasmic tail. Phosphorylation of the ITIM/ITSM by SFKs mediates binding and activation of the Src homology 2 domain-containing protein-tyrosine phosphatases 1 and 2 (Shp1, Shp2). Anti-G6B mAb 17-4 used to generate the G6B-GPVI and G6B-CD32A CAPRIs binds outside the heparan sulfate (HS), heparin ligand binding site of G6B (purple box). CAPRI-mediated co-clustering of G6B with either the ITAM-containing GPVI-FcR γ-chain complex (G6B-GPVI v1 and v2 CAPRIs), or CD32A (FcγRIIA, G6B-CD32A CAPRI) brings the SFK-ITAM-Syk proximal signaling complexes within the molecular reach of G6B ITIM/ITSM-associated Shp1/Shp2. GPVI mAb 3J24 used to generate the G6B-GPVI CAPRI v1 binds outside the ligand-binding site of GPVI (blue box), whereas the VRs of GPVI Fab glenzocimab used to generate the G6B-GPVI v2 CAPRI blocks the ligand binding site of GPVI. CD32A mAb IV.3 used to generate the G6B-CD32A CAPRI blocks to the ligand binding site of CD32A (red box). **(B)** Generation of G6B-GPVI and G6B-CD32A bi-scFv CAPRIs. The light- and heavy-chain variable regions (VRs) of G6B mAb 17-4 were recombinantly joined to the VRs of **(i)** GPVI mAb 3J24, **(ii)** GPVI Fab glenzocimab, **(iii)** CD32A mAb IV.3 using short glycine/serine-rich linker sequences.

Mathematical and structural modelling predicted CAPRIs to be sufficiently short (∼10 nm) to bring G6B-associated Shp1 and Shp2 within the molecular reach of components of the GPVI-FcR γ-chain- and CD32A-ITAM proximal signaling complexes **(Figures 2A and 2B)**. We have measured the molecular length of Shp1 from the N-terminal SH2 domain to the PTP pocket to be 13 nm.(27) Shp2 is predicted to have a similar reach, based on structural similarities. It follows that hetero-clustering G6B with either GPVI or CD32A within approximately this distance would enable G6B-associated Shp1/Shp2 to reach and dephosphorylate tyrosine residues in the cytoplasmic tails of either GPVI or CD32A. Structural model of the ternary complex of G6B, CD32A and G6B-CD32A CAPRI bound to known epitopes on both receptors was constructed by superimposing AlphaFold2-generated structures for complex components **(Figure 2)**. Transmembrane (TM) segments were derived from the Transmembrane Hidden Markov Model (TMHMM). Similar structures and proximities were modelled for G6B, GPVI and G6B-GPVI v1 and v2 CAPRIs, as well as anti-G6B mAb 17.4 bound to G6B and CD32A via its Fab and Fc regions, respectively (*data not shown*). The binding epitope of anti-GPVI mAb 3J24 on GPVI is awaiting experimental confirmation.

**Figure 2.**
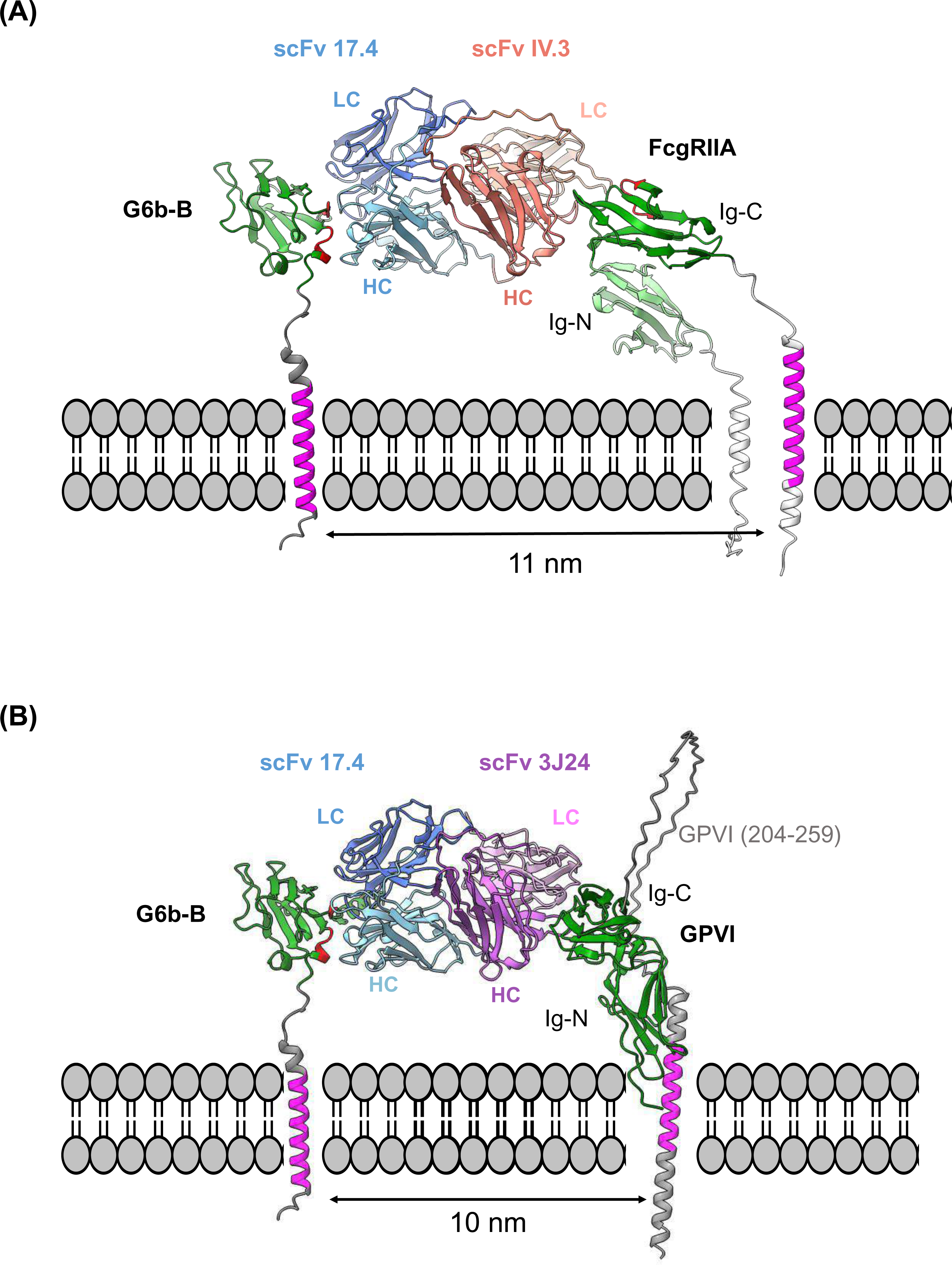
Structural models of ‘receptor bridging complexes’ involving G6b-B, GPVI or FcγRIIA and their CAPRIs. The model was constructed by superimposing AlphaFold2-generated structures for complex components. The models show include scFv fragments specific to G6b-B (17–4), **(A)** FcγRIIA (IV.3) and **(B)** GPVI (3J24), respectively. The construction of the models is described under methods. Approximate distances between the transmembrane regions are indicated. Labels LC, HC indicated the immunoglobulin (Ig) domains of the light and heavy chains, respectively. Transmembrane segments colored in magenta have been derived with Deep TMHMM.(35) DeepTMHMM predicts alpha and beta transmembrane proteins using deep neural networks. https://doi.org/10.1101/2022.04.08.487609) Known epitopes in the ectodomains are colored in red. Sequences C-terminal to the TM helices have been omitted from the view for clarity.

### CAPRIs inhibit ITAM-mediated platelet aggregation

Biological effects of G6B-GPVI and G6B-CD32A CAPRIs were initially investigated by light transmission aggregometry. Pre-treatment of platelets with 10 μg/ml (0.2 μM) G6B-GPVI v1 and v2 CAPRIs inhibited platelet aggregation in response to 3 μg/ml of collagen and 10 μg/ml of CRP **(Figures 3Ai and 3Bi)**, but had no effect on aggregation in response to 3 μg/ml (20 nM) of the anti-CD9 mAb Alma1, which induces CD32A-dependent platelet activation. **(Supplemental Figure 2A)**.(28) The anti-GPVI mAb 3J24 was previously shown to not inhibit GPVI-mediated platelet aggregation.(29) Using an anti-GPVI 3J24 scFv (GPVI scFv) inhibits platelet aggregation **(Figures 3Ai and 3Bi).** However, linking the variable sequences of 3J24 to those of anti-G6B mAb 17.4, as in the G6B-GPVI v1 CAPRI conferred inhibitory activity, comparable to that of glenzocimab **(Figures 3Ai and 3Bi)**. As expected, the G6B-GPVI v2 CAPRI, which shares the same anti-GPVI variable sequences as glenzocimab also inhibited GPVI-induced platelet aggregation **(Figures 3Ai and 3Bi)**. Pretreatment of platelets with 10 μg/ml (0.2 μM) the G6B-CD32A CAPRI inhibited platelet aggregation to anti-CD9 mAb Alma1 **(Figure 3Ci)**, but neither to collagen nor CRP, highlighting the receptor-specificity of these CAPRIs **(Supplemental Figures 3Ai and 3Bi)**. The anti-G6B 17.4 scFv (G6B scFv) had no effect on GPVI- or Alma1-mediated platelet aggregation **(Figure 3Ci)**.

**Figure 3.**
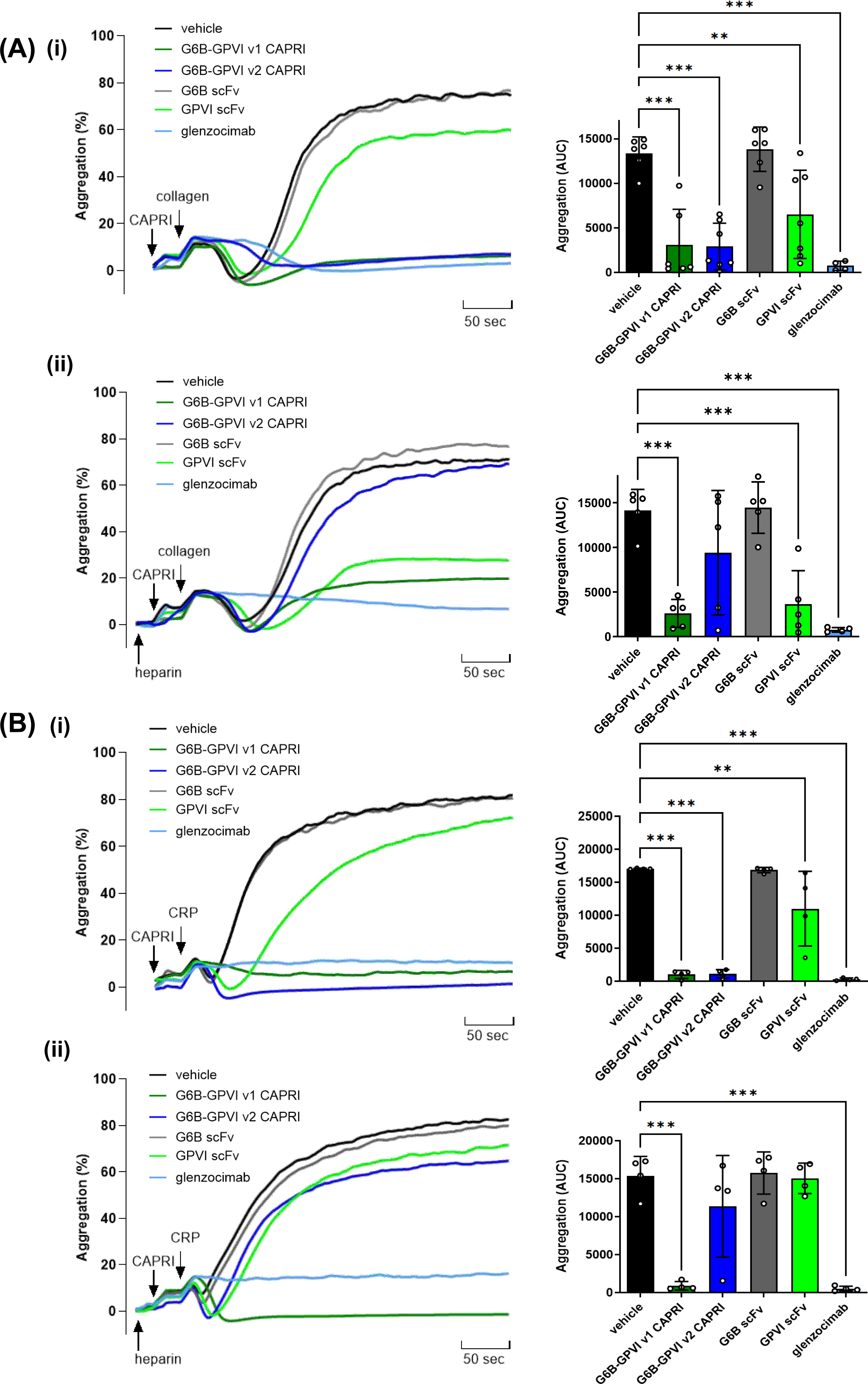
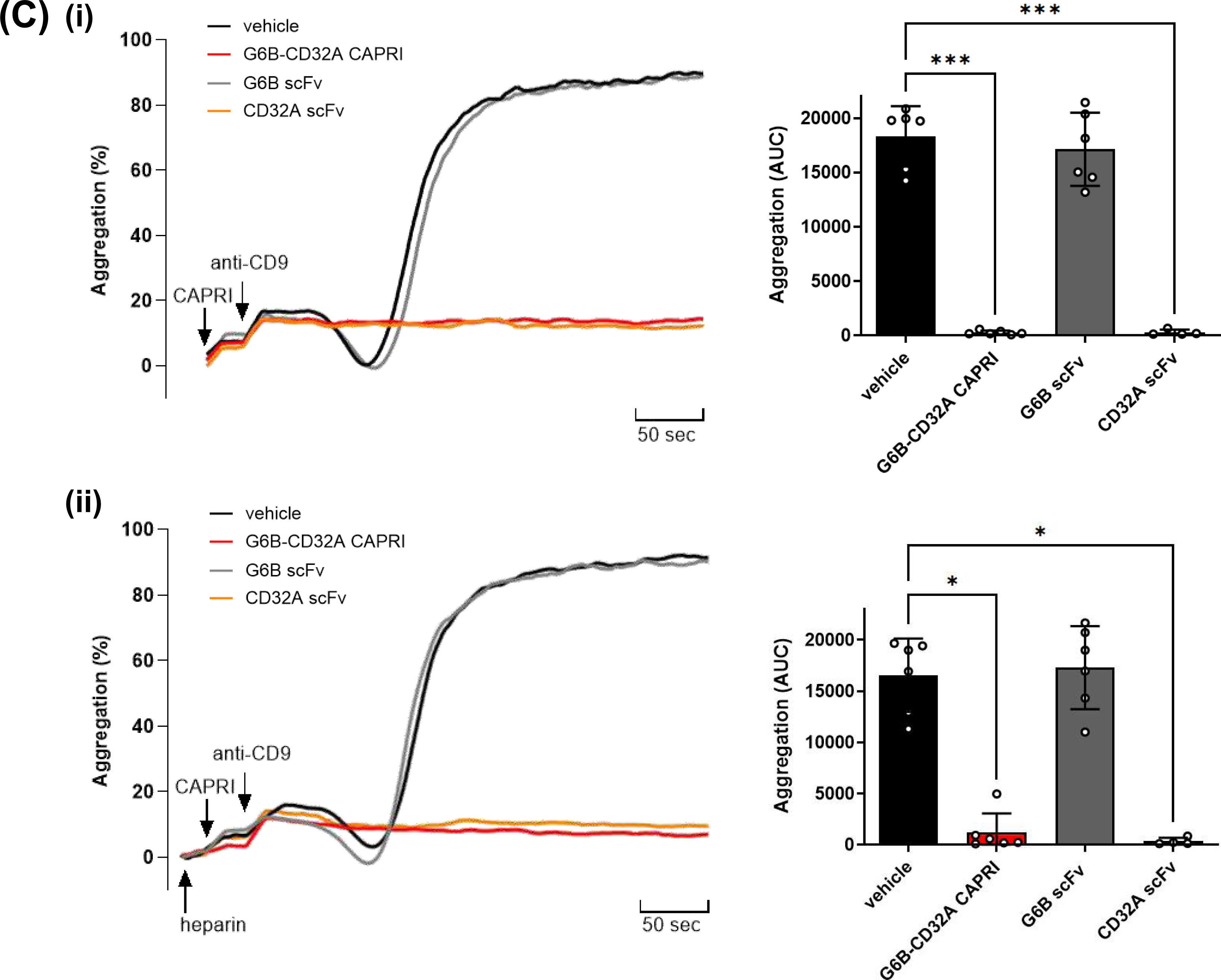
CAPRIs inhibit either GPVI- or CD32A-mediated platelet aggregation. **(A)** Washed human platelets (3 x 10^8^/ml) from healthy donors were pre-treated with either 0.2 μM G6B-GPVI v1, G6B-GPVI v2, G6B-CD32A CAPRIs, G6B scFv, GPVI scFv, CD32A scFv or glenzocimab, in the **(i)** absence and **(ii)** presence of 1 U/ml heparin, prior to stimulation with 3 μg/ml of **(A)** collagen or **(B)** or 10 µg/ml of the GPVI-specific agonist collagen-related peptide (CRP), or **(C)** 3 μg/ml (20 nM) anti-CD9 mAb Alma1. Platelet aggregation was measured in real- time using an APACT 4004 light transmission aggregometer, at 37°C with constant stirring. Representative traces and area under the curve (AUC) are shown. (n = 6 per condition). \**P* < 0.05, \*\**P* < 0.01, \*\*\**P* < 0.001, one-way ANOVA Bonferroni’s test, mean ± SD.

We also investigated the effects of heparin on the inhibitory effects of CAPRIs. Heparin is a ligand of G6B and enhances platelet reactivity to collagen.(9) Platelets were pre-treated with 1 U/ml heparin, prior to addition of either CAPRIs or control compounds, then stimulated with either 3 μg/ml collagen or 10 μg/ml CRP. Under these conditions, G6B-GPVI CAPRI v1 and glenzocimab maintained their inhibitory activities, whereas G6B-GPVI CAPRI v2 did not and GPVI scFv remained ineffective **(Figures 3Aii and 3Bii).** None of them an effect on aggregation in response to 3 μg/ml (20 nM) of the anti-CD9 mAb Alma1 **(Supplemental Figures 2Bi and 2Bii)**. Similarly, both the G6B-CD32A CAPRI and anti-CD32A IV.3 scFv (CD32A scFv) maintained their inhibitory effects on Alma1-mediated platelet aggregation **(Figure 3Cii)**, but did not counteract collagen or CRP **(Supplemental Figures 3Aii and 3Bii)**.

### G6B-CD32A CAPRI inhibits HIT and VITT platelet aggregation

We next investigated the inhibitory activity of the G6B-CD32A CAPRI on HIT- and VITT-induced platelet aggregation. HIT auto-antibodies bind to neo-epitopes exposed on heparin-bound PF4, forming immune complexes that activate platelets via CD32A, the pathological outcome being thrombocytopenia and thrombosis in humans and in mouse models.(30) In contrast, VITT autoantibodies bind to PF4 in the absence of exogenous heparin, triggering CD32A-dependent platelet activation, thrombocytopenia and thrombosis, comparable to HIT immune complexes.(31) Differences in the clinical manifestations of HIT and VITT are thought to be due to the unique biophysical properties of the respective anti-PF4 autoantibodies, the immune complexes that are formed, and their compartmentalization and ability to bind and initiate CD32A signaling.(30)

Consistent with findings using mAb Alma1 to activate platelet CD32A, we found that the G6B-CD32A CAPRI strongly inhibited HIT- and VITT-induced platelet aggregation **(Figures 4A and 4B)**. Two different HIT anti-PF4/heparin antibodies were tested, namely mAb KKO, which is a murine IgG_2b_, and the chimeric antibody 5B9, consisting of a human IgG_1_ Fc and mouse IgG_1_ Fab_2_.(32) Both antibody subclasses bind to human CD32A, but with different affinities.(32, 33) The G6B-CD32A CAPRI also inhibited platelet aggregation to the VITT anti-PF4 chimeric antibody 1E12, which has a human IgG_1_ Fc and murine Fab_2_ **(Figures 4B and 4Cii)**. The G6B scFv had no effect on either the HIT antibody 5B9- or VITT antibody 1E12-mediate platelet aggregation or shape change **(Figures 4A and 4B)**. Neither the G6B-GPVI CAPRIs nor control scFvs inhibited either HIT- or VITT-induced platelet aggregation **(Supplemental Figures 4A and 4B)**.

**Figure 4.**
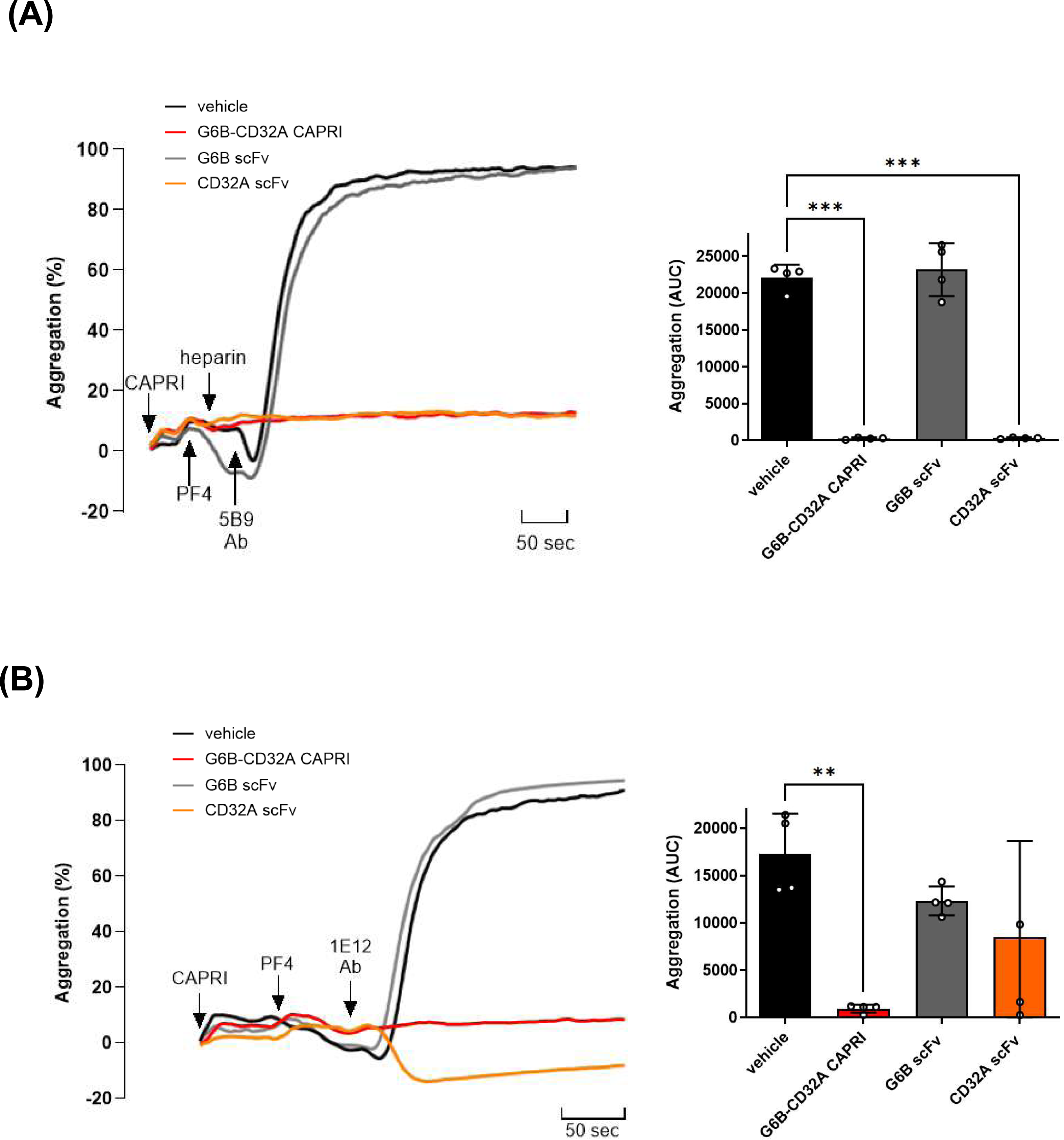
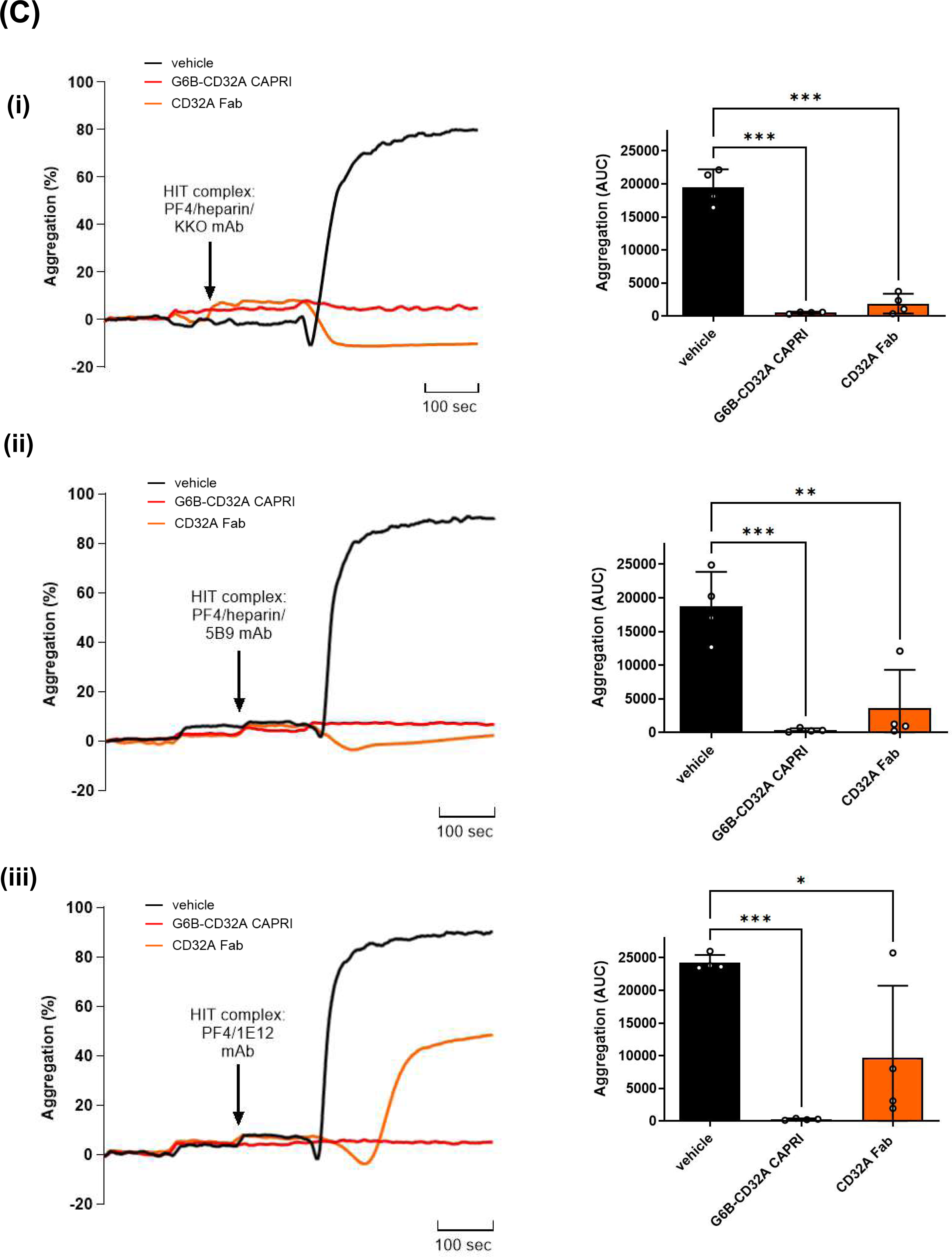
G6B-CD32A CAPRI inhibits immune complex-mediated platelet aggregation. **(A)** Washed human platelets (3 x 10^8^/ml) from healthy donors were pre-treated with either 0.2 μM G6B-CD32A CAPRIs, G6B and CD32A scFvs prior stimulation. Platelets were stimulated with a **(i)** HIT immune complex consisting of PF4/heparin/mAb KKO by sequentially adding 10 μg/ml PF4, 0.5 U/ml heparin and 100 μg/ml KKO at 30-seconds intervals to platelet suspensions after adding 1 mg/ml fibrinogen. **(ii)** Same as in (i), but 50 μg/ml mAb 5B9 was used rather than KKO to stimulate platelets. **(B)** Same as in (i), but platelets were stimulated with the VITT mAb 1E12. **(C)** Improved efficacy of G6B-CD32A CAPRI compared with CD32A Fab. Washed human platelets were treated with either 0.2 μM G6B-CD32A CAPRI or CD32A Fab prior to stimulation with either **(i)** KKO, or **(ii)** 5B9 HIT, **(iii)** or VITT 1E12. Representative traces and area under the curve (AUC) are shown. (n = 4-6 per condition). \**P* < 0.05, \*\**P* < 0.01, \*\*\**P* < 0.001, one-way ANOVA Bonferroni’s test; mean ± SD.

We also compared the inhibitory activities of G6B-CD32A CAPRI and anti-CD32A IV.3 Fab, which share the same anti-CD32A variable sequences. The CD32A Fab inhibited HIT-induced platelet aggregation, but not shape change, whereas the G6B-CD32A CAPRI inhibited both shape change and aggregation **(Figure 4Ci and 4Cii)**. Similarly, the CD32A Fab was less effective at inhibiting VITT-induced platelet aggregation compared with the G6B-CD32A CAPRI, which inhibited both shape change and aggregation **(Figure 4Ciii)**. Findings demonstrate improved efficacy of the CAPRI compared with the monovalent CD32A Fab.

### CAPRIs inhibit of GPVI and CD32A proximal signaling

We next investigated the inhibitory effects of CAPRIs on GPVI and CD32A proximal signaling using a capillary-based immunoassay to measure markers of activation. Platelets were pretreated with either 0.2 μM CAPRIs or control compounds before stimulation with either 30 μg/ml CRP or 10 μg/ml (60 nM) anti-CD9 mAb Alma1. Phosphorylation of Syk tyrosine residues 525/526 (p-Tyr525/526) is an indirect indicator of Syk activation, and one of the earliest and sensitive markers of initiation of GPVI and CD32A signaling.(10) We demonstrate that G6B-GPVI v1 and v2 CAPRIs specifically inhibit Syk p-Tyr525/526 downstream of GPVI **(Figures 5Ai and 5Aii),** whereas G6B-CD32A CAPRI did not, demonstrating receptor specificity. Both G6B-GPVI CAPRIs also inhibited GPVI-induced Shp1 p-Tyr564 and Shp2 p-Tyr542, suggesting regulation of function, as well as SFK p-Tyr418 phosphorylation **(Figures 5Ai and 5Aii)**, an indirect indicator of tyrosine kinase activity essential for ITAM phosphorylation.(10) Conversely, G6B-CD32A CAPRI specifically inhibited Syk p-Tyr525/526, Shp1 p-Tyr564 and Shp2 p-Tyr542 downstream of CD32A, whereas G6B-GPVI v1 and v2 CAPRIs did not **(Figures 5Bi and 5Bii).** Glenzocimab had a similar effect on GPVI signaling as the G6B-GPVI v1 and v2 CAPRIs, whereas the GPVI scFv had only a marginal effect **(Figures 5Ai and 5Aii)**, and the CD32A Fab had a similar effect to G6B-CD32A CAPRI on CD32A signaling. The G6B scFv had no effect on either GPVI or CD32A signaling, suggesting that the mode of action of CAPRIs is by hetero-clustering the co-inhibitory receptor G6B with either of the ITAM receptors.

**Figure 5.**
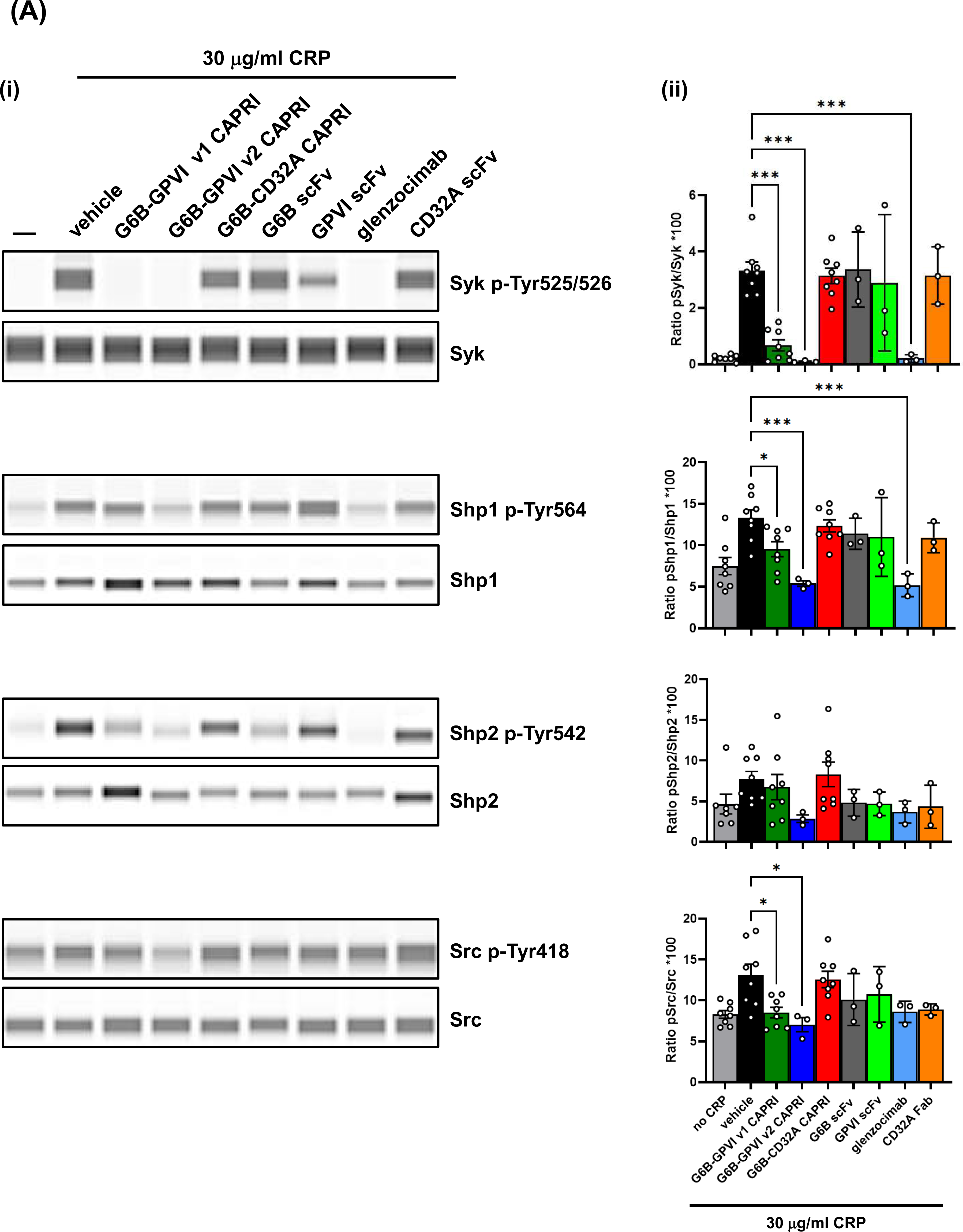
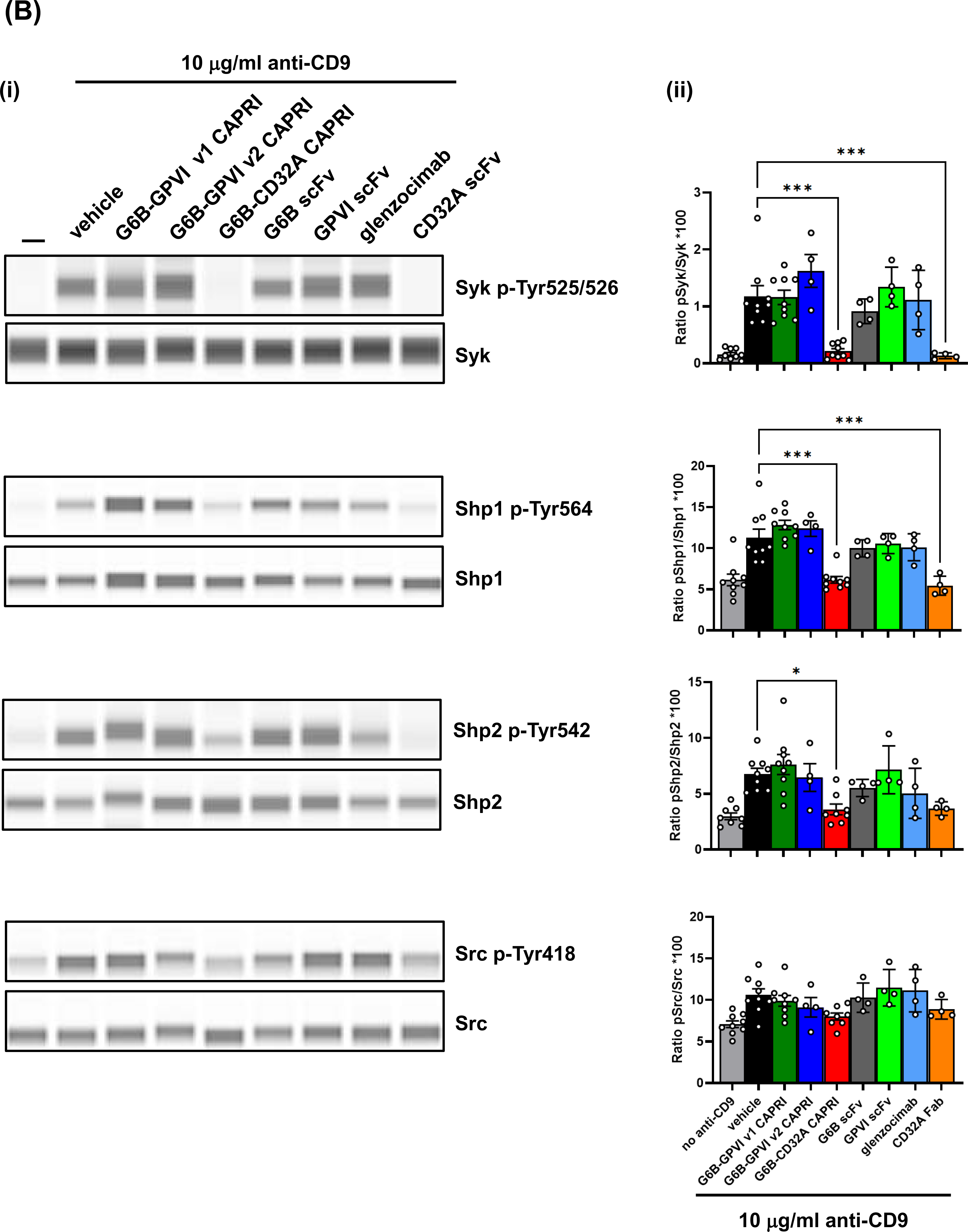
CAPRIs inhibit GPVI and CD32A proximal signaling. Washed human platelets (5 × 10^8^/ml) were incubated with 0.2 μM G6B-GPVI v1, G6B-GPVI v2, G6B-CD32A CAPRIs, G6B scFv, GPVI scFv, CD32A scFv or glenzocimab for 15 minutes at 37°C prior to stimulation with **(A)** 30 μg/ml CRP or **(B)** 10 μg/ml anti-CD9 mAb Alma1 for 90 seconds before lysing and investigation of named phospho-tyrosine sites. **(i)** Representative blots of capillary-based immunoassays with the indicated antibodies **(ii)** and quantification of peak areas are shown. n=4-8 experiments/conditions. Asterisks refer to significant difference compared with control samples within conditions. *P<.05, **P< .01, ***P < .001, Krustal-Wallis’s test; mean ± SD.

### CAPRI inhibition of platelet aggregation on thrombogenic surfaces

We next investigated the inhibitory effects of the CAPRIs on platelet adhesion and aggregation on collagen under arterial shear. Predictably, 10 μg/ml (0.2 μM) of either G6B-GPVI v1 or v2 CAPRIs significantly reduced platelet adhesion, aggregation and thrombus growth of anticoagulated whole blood flowed at 1,500 s^−1^ over a collagen-coated surface, compared with G6B-CD32A CAPRI, G6B and CD32A scFvs, which had no effect **(Figures 6Ai and 6Aii, 6Bi and 6Bii and Supplemental Videos 1-8)**. Glenzocimab was as effective as G6B-GPVI v1 CAPRI at inhibiting platelet adhesion, aggregation and thrombus growth under these conditions, whereas the GPVI scFv had a partial effect. The G6B-GPVI v2 CAPRI was most effective at inhibiting platelets to adhere and form aggregates, as well as reducing thrombus growth on collagen under these conditions, suggesting that blocking both GPVI ligand binding and downstream signaling is more efficacious then blocking either alone **(Figures 6Ai and 6Aii, 6Bi and 6Bii and Supplemental Videos 1-8).**

**Figure 6.**
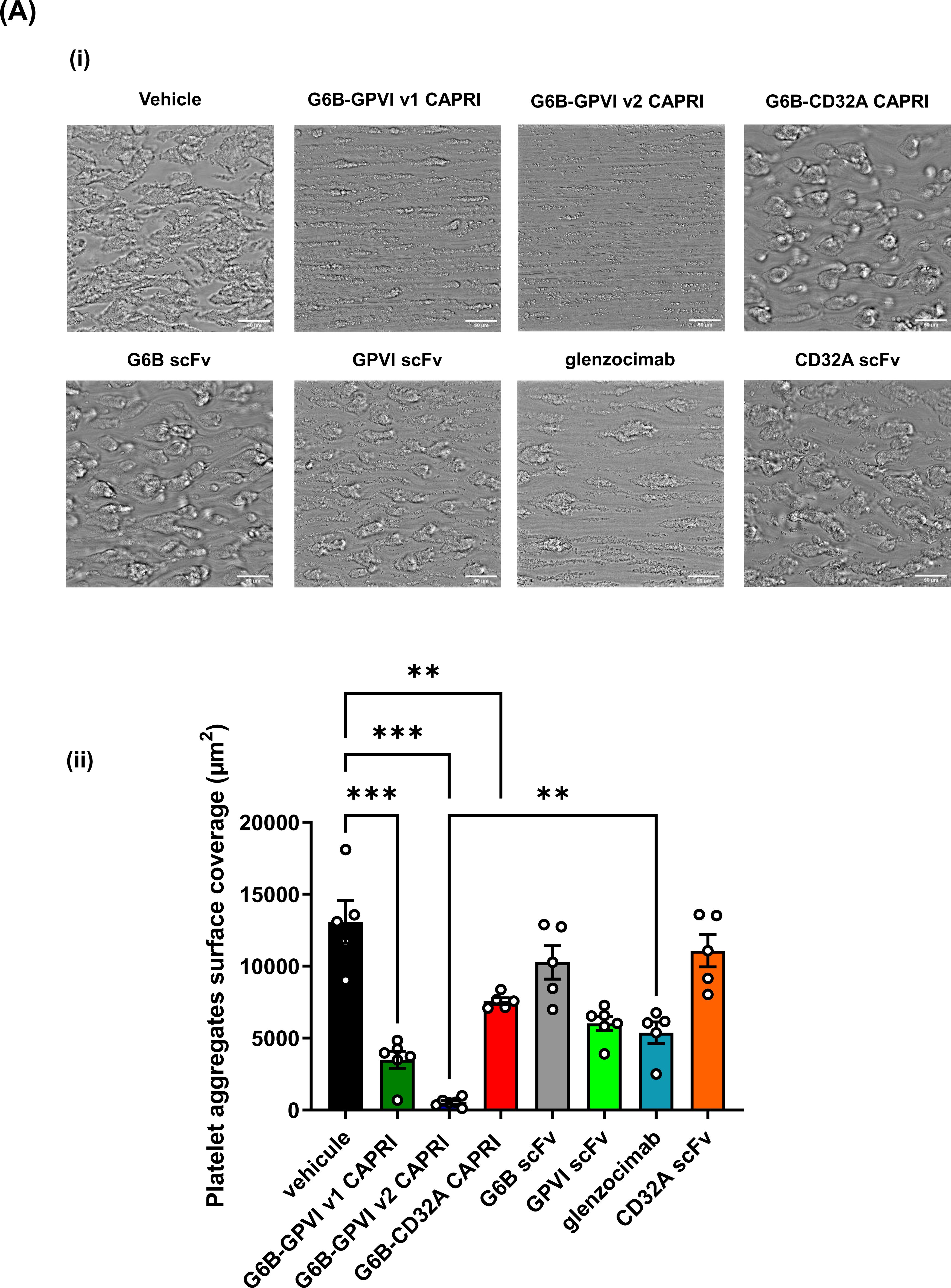
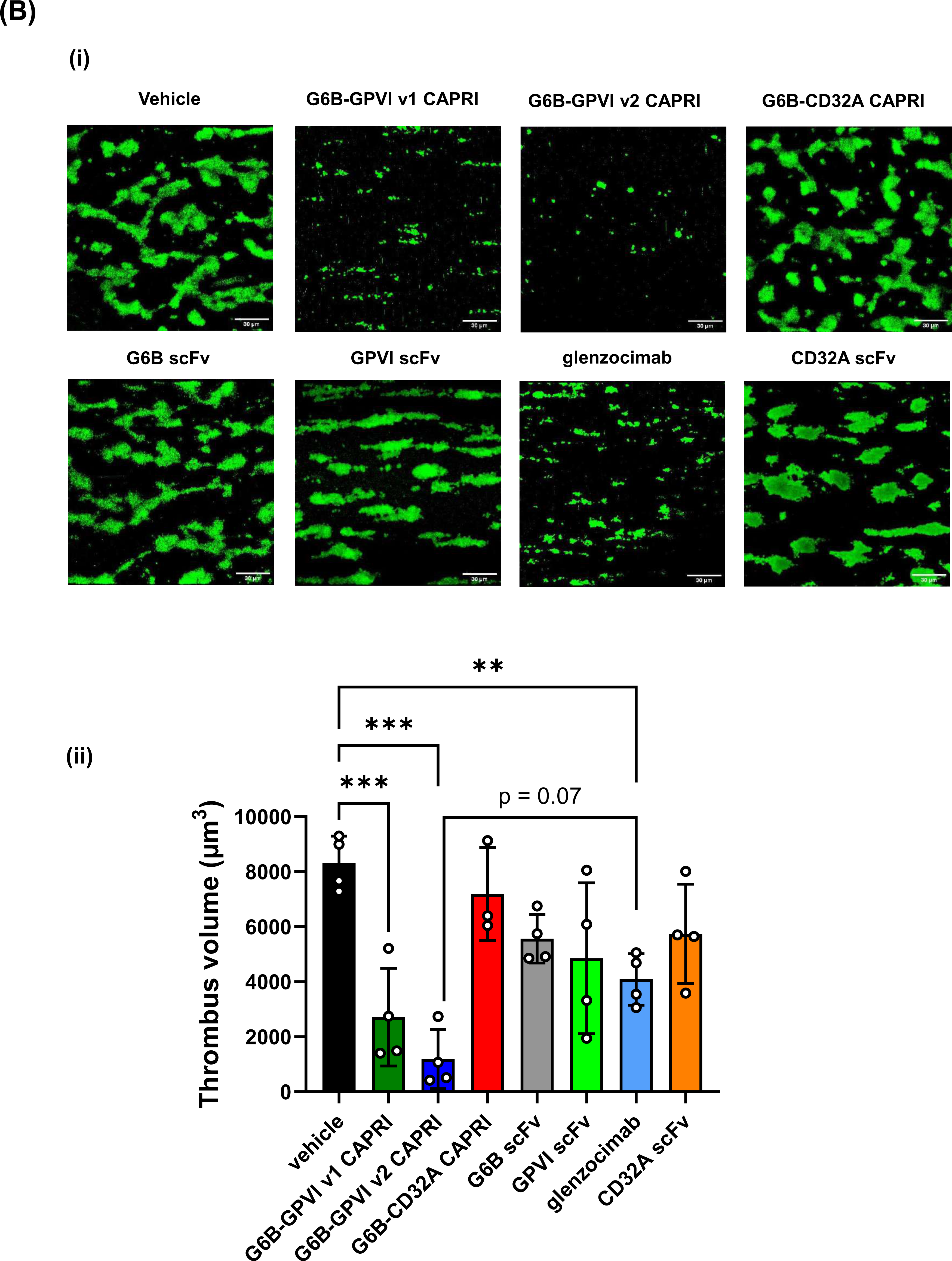
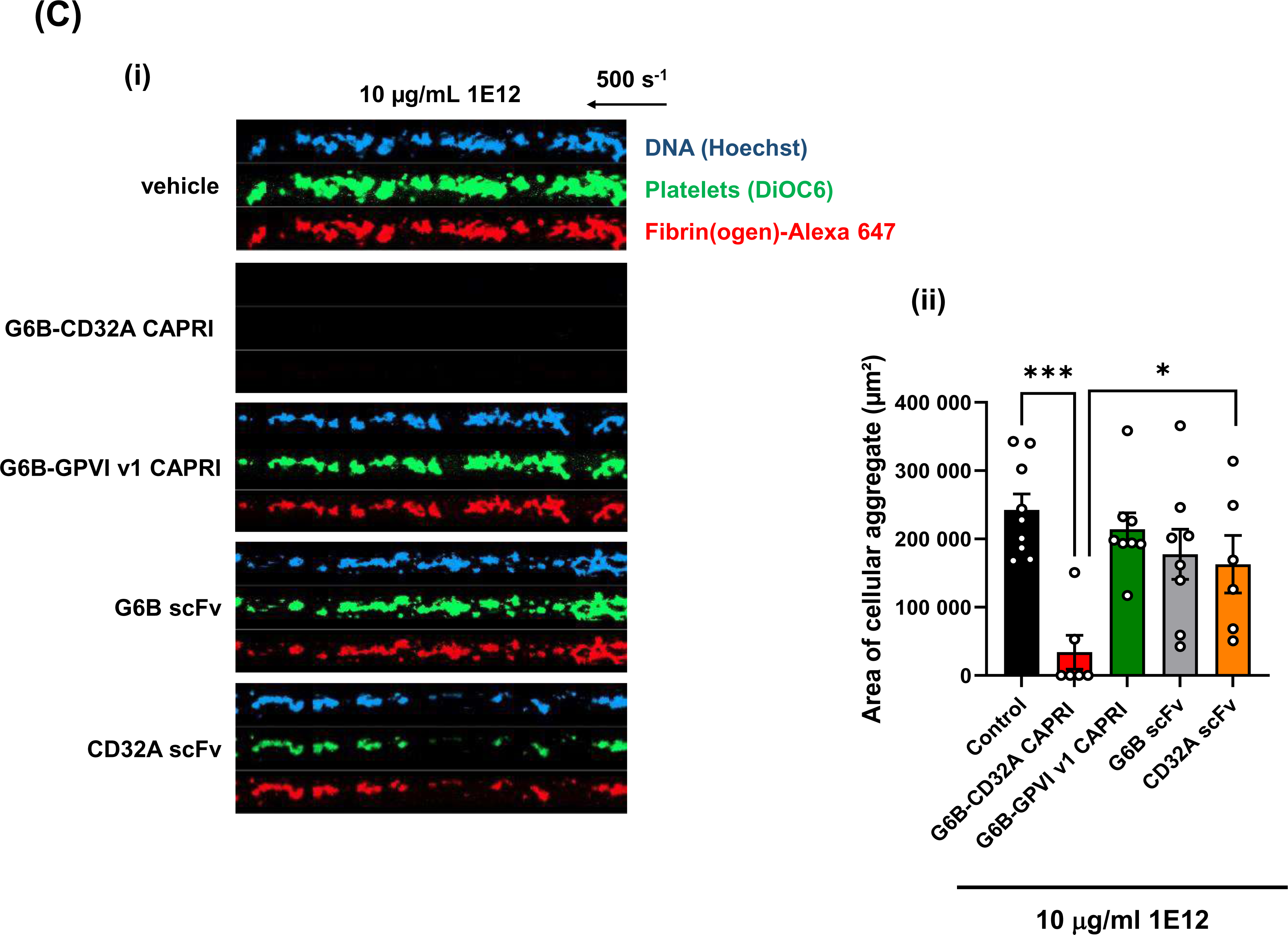
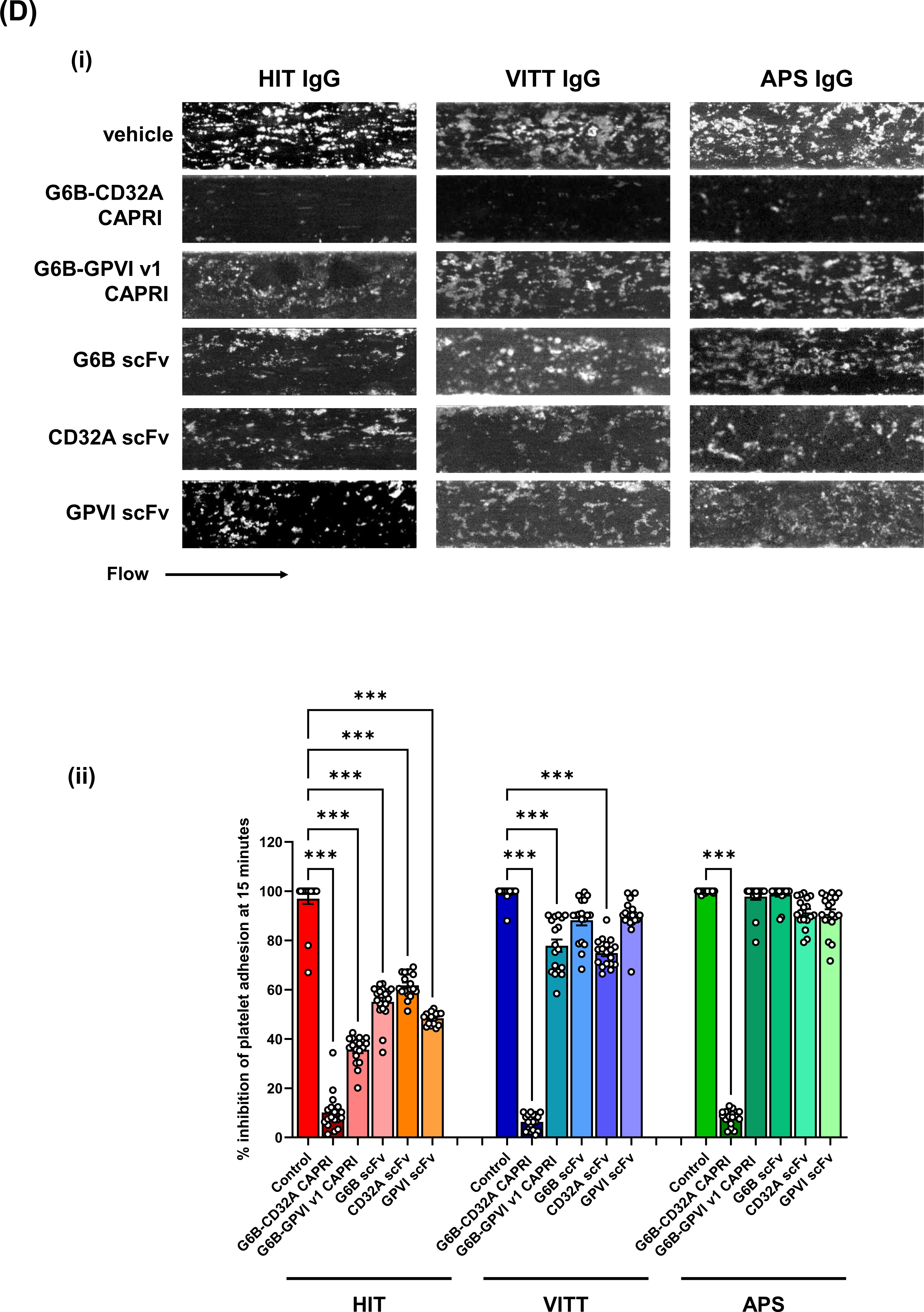
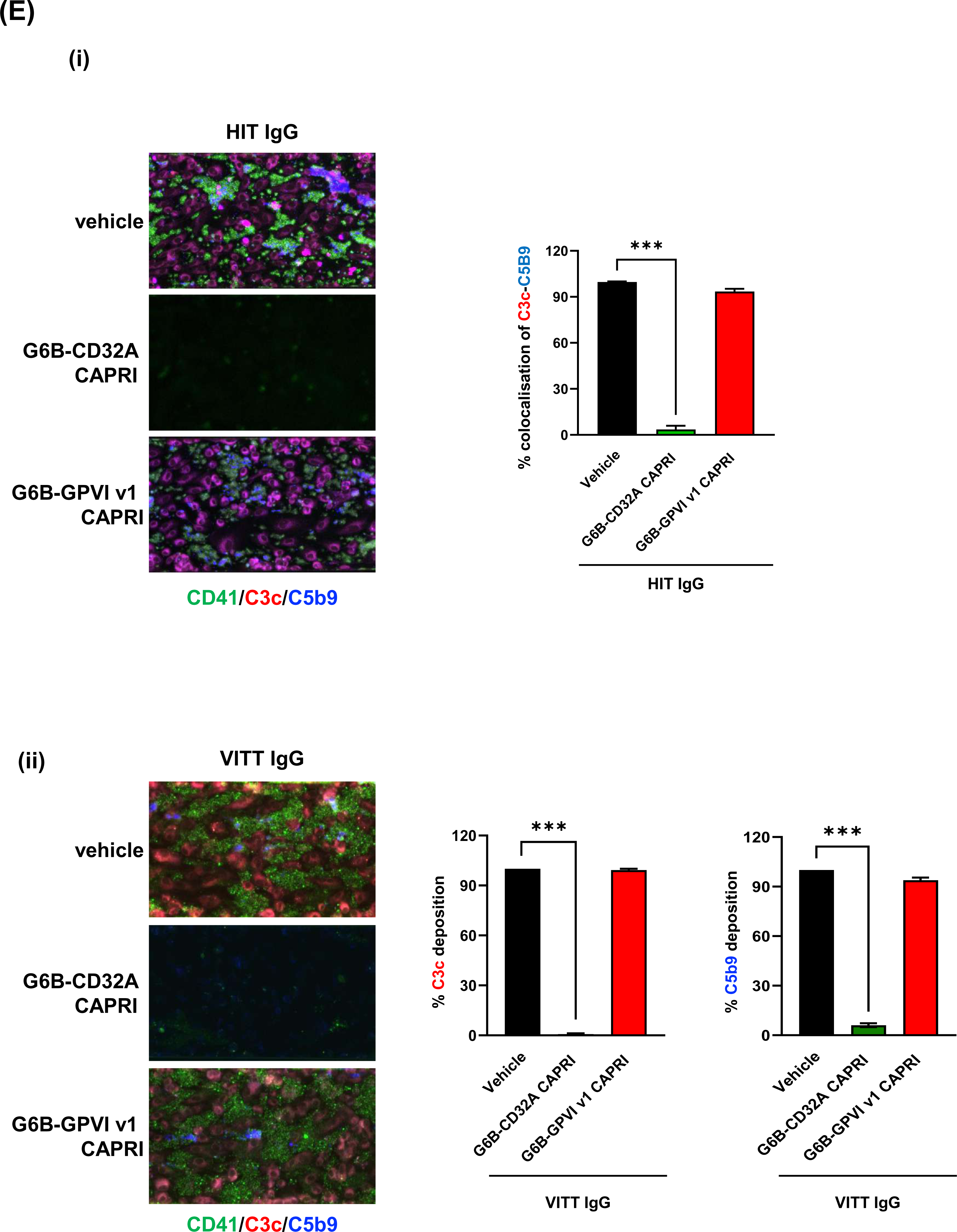
CAPRIs inhibit thrombus formation on thrombogenic surfaces. **(A)** Whole, hirudinated blood collected from healthy donors was treated with 0.2 μM of G6B-CD32A, G6B-GPVI v1 or G6B-GPVI v2 CAPRIs, G6B scFv, GPVI scFv, CD32A scFv, or glenzocimab for 15 minutes at 37°C prior to being flowed over a surface coated with 200 μg/ml collagen at 1,500 s^-1^ for 5 minutes at 37°C. **(i)** Representative DIC images of adherent platelets on collagen fibrils were captured, scale bars represent 50 µm. **(ii)** Percentage of surface coverage was quantified. Data presented are means ± SD of 5 independent experiments. Data was analyzed by one-way Anova and post hoc Turkey’s test (*\*P*<0.01, *\*\*P*<0.01, \**\*\*P*<0.01). **(B)** Whole, hirudinated blood collected from healthy donors was labelled with DIOC_6_ and treated with 0.2 μM of G6B-CD32A, G6B-GPVI v1 or G6B-GPVI v2 CAPRIs, G6B scFv, GPVI scFv, CD32A scFv, or glenzocimab for 15 minutes at 37°C prior to being flowed over a surface coated with 200 μg/ml collagen at 1,500 s^-1^ for 5 minutes at 37°C. **(i)** Platelet aggregation was visualized in random fields by confocal microscopy, scale bars represent 50 µm. **(ii)** The volume of the thrombi was quantified. Data presented are means ± standard deviation of 4 independent experiments. Data was analyzed by one-way Anova and post hoc Turkey’s test (*\*P*<0.01, *\*\*P*<0.01, \**\*\*P*<0.01). **(C)** Recalcified whole blood from healthy donors was treated with 0.2 μM of G6B-CD32A CAPRI, G6B-GPVI v1 CAPRI, G6B scFv or CD32A scFv, then 6 minutes with 10 μg/mL 1E12 prior to being flowed over a VWF-coated microfluidic channels. **(i)** Images corresponding to areas of 0.1 mm^2^ are shown with platelets in green (DiOC_6_), fibrin(ogen) in red (Alexa Fluor 647-labeled fibrinogen), and leukocytes in blue (Hoechst 33342, DNA dye). **(ii)** The mean areas covered by platelets were calculated for each condition (n = 6 independent experiments), by measuring using ImageJ software the surface covered by large aggregates (> 100 μm^2^) in 20 different areas. Data was analyzed by one-way Anova and post-hoc Turkey’s test (*\*P*<0.01, *\*\*P*<0.01, \**\*\*P*<0.01). **(D)** Effect of CAPRIs on HIT, VITT and APS thrombosis in a microfluidic system wherein the channels were coated with near-confluent HUVECs on fibronectin. **(i)** Representative images over a 15-minute window of flow through the channels from 1 of 3 independent studies showing platelet accumulation (white aggregates) on the endothelial lining. Scale bar is included, and arrows at the bottom indicate direction of flow. In this assay, optimal platelet adhesion occurs HUVECs-lined channels subjected to green laser-photochemical injury by hematoporphyrin,(18) followed by perfusion of whole blood supplemented with either PF4 (25 μg/ml) and HIT IgG (100 μg/ml), VITT IgG (100 μg/ml) or APS IgG (100 μg/ml) along with 0.6 μM of G6B-CD32A, G6B-GPVI v1 CAPRIs or G6B scFv, CD32A scFv, GPVI scFv. **(ii)** Quantitative analysis of platelet adhesion showing mean ± SEM of 3 independent experiments. Data was analyzed by one-way Anova and post-hoc Turkey’s test (*\*P*<0.01, *\*\*P*<0.01, \**\*\*P*<0.01). (**E)** Effect of CAPRIs on complement activation in the same microfluidic system as in (D). (**i)** Representative confocal Z stack images of anti-human C3c and anti-human C5b9. Injured channels were fixed and stained after 15 minutes of pre-mixed blood flow with HIT IgGs along with CAPRIs (0.6 μM). Cells were fixed and stained for 1 of 3 independent studies showing platelet accumulation (green aggregates) on the endothelial lining. (**ii)** Quantitative analysis of % colocalized of complement component from 3 independent studies in triplicate expressed in mean ± SEM are shown. n=3 independent experiments. Data was analyzed by one-way Anova and post-hoc Turkey’s test (*\*P*<0.01, *\*\*P*<0.01, \**\*\*P*<0.01).

We next tested the effects of the CAPRIs on VITT-induced thrombus formation on VWF under flow. Whole blood from healthy donors was incubated with VITT antibody 1E12 and perfused over a VWF-coated surface under venous shear (500 s^−1^). Numerous large fibrin(ogen)-containing platelet/leukocyte aggregates were observed in the absence of CAPRIs or controls **(Figure 6Ci)**, as previously reported.(17) Consistent with findings from platelet aggregation assays in suspension, thrombus formation induced by 1E12 was completely inhibited with 0.2 μM G6B-CD32A CAPRI, whereas the G6B-GPVI v1 CAPRI and G6B scFv had no effect **(Figures 6Ci and 6Cii)**. The CD32A scFv partially inhibited thrombus formation under conditions tested, demonstrating improved efficacy of the G6B-CD32A CAPRI, which blocks both ligand binding and downstream signaling.

### CAPRI inhibition of immune-induced thrombus formation on injured endothelial cells

We further assessed the ability of CAPRIs to block thrombus formation in an endothelial cell-coated microfluidic assay mimicking the immune-induced prothrombotic disorders HIT, VITT and APS, as previously described **(Supplemental Figure 5A)**.(18) In this assay, thrombus formation occurs following photochemical injury of endothelial cells and perfusion of whole blood pretreated with HIT, VITT or APS immune complexes, consisting of anti-PF4/HIT, -PF4/VITT or -APS IgG antibodies, respectively. Addition of G6B-CD32A CAPRI to the blood 15 minutes following addition of HIT, VITT or APS immune complexes and photochemical injury markedly reduced platelet accumulation on the injured endothelium **(Figures 6Di and 6Dii and Supplemental Figures 6A-C)**. The G6B-GPVI v1 CAPRI and scFv controls had marginal or no significant effects at inhibiting thrombus formation under these conditions **(Figures 6Di and 6Dii and Supplemental Figures 6A-C).** Furthermore, G6B-CD32A CAPRI also decreased HIT- and VITT-induced complement activation, as shown by the inhibition of deposition of the early and terminal components of the complement cascade C3c and C5b9, respectively on platelets **(Figures 6Ei and 6Eii).** Collectively, these HUVEC microfluidic findings demonstrate that the G6B-CD32A CAPRI is highly effective at inhibiting HIT, VITT and APS immune complex-dependent thrombus formation on injured endothelial cells under flow.

## Discussion

Findings from this study further demonstrate the importance of hetero-clustering ITIM and ITAM receptors in regulating platelet activation. We provide multifaceted evidence to establish CAPRIs as efficacious ITAM receptor-specific inhibitors of platelet activation and demonstrate distinct advantages of CAPRIs over Fab fragments that block a single ITAM receptor, namely either GPVI or CD32A. Prototype CAPRIs provide proof-of-concept of harnessing physiological co-inhibitory receptors to attenuate cellular reactivity to pathophysiological factors, irrespective of receptor-ligand engagement, a form of “personalized platelet medication”. CAPRIs also provide a means of augmenting the efficacy of antibody-based therapies targeting platelet activation receptors and potentially reducing dosing regiments while enhancing platelet-targeting.

A primary objective of translational platelet research is decoupling platelets’ thrombotic activity from their hemostatic capacity. Despite many studies claiming to have identified putative targets or compounds that address this issue, all current anti-platelet therapies cause increased bleeding outcomes in patients, questioning whether decoupling is attainable. The commonality of all current anti-thrombotic approaches that may underlie this shortcoming is their blockade of vital platelet activation and adhesion receptors, and associated signaling pathways. To achieve decoupling, we devised an alternative strategy utilizing the physiological co-inhibitory receptor G6B to drive platelets towards less active states, irrespective whether in suspension or ligand-bound at sites of injury **(Figures 7A and 7B)**. Cross-linking of platelet ITIM and ITAM receptors could provide treatments for a number of diseases and conditions where platelet activation is an integral component, such as HIT- or VITT-associated thrombosis.

**Figure 7.**
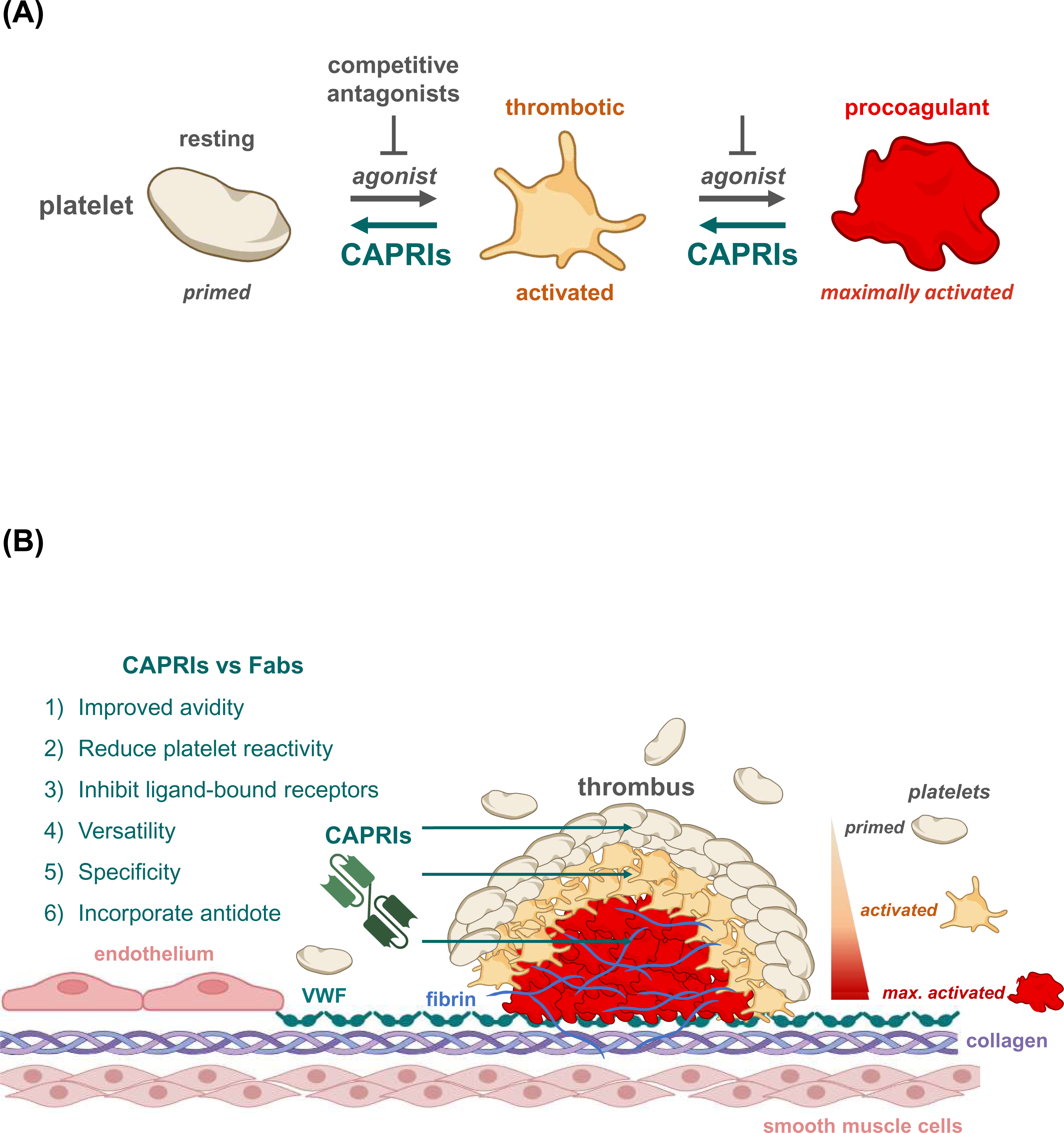
Model of how CAPRIs provide a paradigm shift in targeting platelets and thrombosis. **(A)** Schematic of CAPRIs shifting the balance of platelets from right to left towards a more quiescent state. CAPRIs offer a versatile and refined approach of regulating the threshold of platelet activation, irrespective if platelet are ligand-bound at sites of injury or circulating in a resting state. This strategy of harnessing a physiological inhibitory of platelet activation goes beyond blocking platelet binding to thrombogenic substrates and positive feedback pathways. **(B)** Schematic depicting several of the potential advantages of CAPRIs compared with conventional anti-thrombotic therapies targeting platelet activation receptors and associated signaling pathways, possibly leading to improved antithrombotic, but with fewer bleeding outcomes in patients.

There are several distinct advantages of bispecific CAPRIs compared with monospecific Fab antagonists **(Figure 7B)**. Apart from improved avidity, CAPRIs are more versatile than Fabs, because they can be engineered to inhibit signaling from unbound and ligand-bound ITAM receptors. There is compelling evidence that low-level tonic signaling from unbound ITAM receptors is required for developmental and cellular responses of hematopoietic cells, the extent and duration of which is dependent on ITAM accessibility and the opposing activities of SFKs and PTPs to phosphorylate and dephosphorylate, respectively. CAPRIs can eliminate this tonic signaling by juxtaposing p-ITIM-associated Shp1 and Shp2 with proximal signaling components of ITAM receptors that can be dephosphorylated and reducing platelet reactivity.

Inhibiting signaling from ligand-bound receptors may be advantageous at inactivating platelets and disrupting preformed thrombi, either alone or in combination with “clot busters”, such as recombinant tissue plasminogen activator (rtPA). Indeed, the G6B-GPVI v1 CAPRI, which binds outside the ligand binding sites of both G6B and GPVI, exhibited potent inhibitory activity of GPVI-mediated platelet activation and function, whereas the monovalent scFvs had no significant activity. In contrast, the G6B-GPVI v2 CAPRI, which shares the same variable sequences as glenzocimab inhibits both ligand binding and downstream signaling, potentially explaining why it was more effective at inhibiting platelet thrombus formation on collagen under flow, compared with G6B-GPVI v1 CAPRI and glenzocimab. Other contributing factors are improved affinity and avidity of the G6B-GPVI v2 CAPRI. Similarly, the G6B-CD32A CAPRI, which shares the same variable sequences as the anti-CD32A ligand blocking antibody IV.3 was also more effective than either the IV.3 Fab or scFv at inhibiting CD32A-mediated platelet activation and function in some instances, notably VITT immune complex-mediated platelet activation.

A hypothetical risk of CAPRIs is that they cause platelet agglutination through *trans*-interactions of receptors on different platelets, as is the case for the bispecific T cell engager (BiTE) blinatumomab, which binds CD3 on T cells and CD19 on B cell lymphoma cells.(34) However, this was not observed for any of the CAPRIs or conditions used in this study, presumably because the probability of CAPRIs engaging two receptors on the same platelet surface is higher than *trans-* interactions with receptors on separate platelets. The possibility of *trans-*interactions occurring at higher concentrations of CAPRIs and platelets remains plausible. Further, we do not anticipate CAPRIs as those described in this study impacting MK development, maturation or platelet production, but will depend on therapeutic windows and dosing regiments that will be determined as part of future *in vivo* testing, requiring animal models expressing humanized receptors on their MKs and platelets, or xenotransplanted with human hematopoietic cells.

Collectively, these findings highlight the superior versatility of CAPRIs compared with Fabs that can be exploited to improve efficacy by swapping and affinity tuning variable sequences to optimize binding properties, varying the length and composition of linker peptide sequences, and modulating the valence of CAPRIs to enhance accessibility and stoichiometry of receptor clustering. We also envisage cleavage sites being embedded within linker peptides, allowing ITIM and ITAM receptors to be uncoupled in the presence of an exogenous catalyst and acting as an “antidote” to restore platelet reactivity to reverse excessive bleeding.

## Acknowledgements

This study was supported by Inserm, the Agence Nationale pour la Recherche (YAS, ANR-23-TARGITT-CE18-0020-01), Etablissement Français du Sang Grand-Est and the National Institute of Health (MP, R35 HL150698).

## Author Contributions

A.M. Designed, performed experiments, analyzed data, wrote and revised the manuscript.

O.B. Performed experiments, analyzed data, revised the manuscript.

A.S. Performed experiments, analyzed data, revised the manuscript.

J.A. Performed experiments, analyzed data, revised the manuscript.

A.B. Performed experiments, analyzed data and revised the manuscript.

C.L. Performed experiments, analyzed data and revised the manuscript.

F.J. Provided VR sequences and reagents, discussed and revised the manuscript.

J.W. Designed, produced CAPRIs, discussed and revised the manuscript.

M.A. Designed, produced CAPRIs, discussed and revised the manuscript.

O.D. Mathematical modelling, discussed and revised the manuscript.

C.V. Designed experiments, analyzed data and revised the manuscript.

L.R. Designed experiments, analyzed data and revised the manuscript.

K.F. Structural modelling, discussed and revised the manuscript.

J.R. Provided reagents, designed experiments, discussed and revised the manuscript.

M.P. Provided reagents, designed experiments, analyzed data and revised the manuscript.

Y.A.S. Conceptualized, designed experiments, analyzed data, wrote and revised the manuscript.

## Data Availability Statement

The data that support the findings of this study are available from the corresponding author upon reasonable request.

