## Supplemental Figures for "G6b-B antibody-based *cis*-acting platelet receptor inhibitors (CAPRIs) as a new family of anti-thrombotic therapeutics"

### Supplemental Figure 1

#### (A) G6B-GPVI version 1 bispecific single-chain variable fragment

**Monoclonal antibody 17-4 $V_L$-linker-$V_H$-linker-Monoclonal antibody 3J24 $V_L$-linker-$V_L$-Histidine (H) tag**

**17-4 VL**

DIQMTQTTSSLSASLGDRVTISCRASQDISNYLNWYQQKPDGTVKLLIYYTSTLHSGVPSRFSGSGSGTDYSLTISNLEQEDVATYFCQ

**Linker** **17-4 VH**

QGYTLPWTFGGGTKLEIKGGGGSGGGGSGGGGSEVQLQQSGAELVKPGASVKLSCTASGFNIKETYIHWVKQRPEQGLEWIGRID

**Linker** **3J24 VH**

CARSYGSSYGIDYWGQGTSVTVSSGGGGSEIQLQQSGPELKKPGETVKISCKASGYTFTNYGMNWVKQAPGKGLKWMGWLNTY

**Linker**

TGESIYPDDFKGRFAFSSETSASTAYLQINNLKNEDMATYFCARGDYGYDDPLDYWGQGTSVTVSSGGGGSGGGGSGGGGSDVV

**3J24 VL**

MTQTPPSVPVTPGESVSISCRSSKSLLHTNGNTYLHWFLQRPGQSPQLLIYRMSVLASGVPDRFSGSGSGTAFTLSISRVEAEDVGVF

**His-Tag**

YCMQHLEYPLTFGAGTKLELK**HHHHHH**

#### (B) G6B-GPVI version 2 bispecific single-chain variable fragment

**Monoclonal antibody 17-4 $V_L$-linker-$V_H$-linker-Monoclonal antibody Glenzocimab $V_H$-linker-$V_L$-Histidine (H) tag**

**17-4 VL**

DIQMTQTTSSLSASLGDRVTISCRASQDISNYLNWYQQKPDGTVKLLIYYTSTLHSGVPSRFSGSGSGTDYSLTISNLEQEDVATYFCQ

**Linker** **17-4 VH**

QGYTLPWTFGGGTKLEIKGGGGSGGGGSGGGGSEVQLQQSGAELVKPGASVKLSCTASGFNIKETYIHWVKQRPEQGLEWIGRID

**Linker**

PADVYGRYDPKFQGKATITADTSSNSAYLQVSSLTSEDTAVYYCARSYGSSYGIDYWGQGTSVTVSSGGGGSQVQLVQSGAEVKKP

**Glenzocimab VH**

GASVKVSCKASGYTFTSYNMHWVRQAPGQGLEWMGGIYPGNGDTSYNQKFQGRVTMTRDTSTSTVYMELSSLRSEDTAVYYCA

**Linker** **Glenzocimab VL**

RGTVVGDWYFDVWGQGTLVTVSSGGGGSGGGGSGGGGSDIQMTQSPSSLSASVGDRVTITCRSSQSLENSNGNTYLNWYQQKP

**His-Tag**

GKAPKLLIYRVSNRFSGVPSRFSGSGSGTDFTFTISSLQPEDIATYYCLQLTHVPWTFGQGTKVEITRHHHHHH

### Supplemental Figure 1

#### (C) G6B-CD32A bispecific single-chain variable fragment

**Monoclonal antibody 17-4 $V_L$-linker-$V_H$-linker-Monoclonal antibody IV.3 $V_H$-linker-$V_L$-Histidine (H) tag**

**17-4 VL**

DIQMTQTTSSLSASLGDRVTISCRASQDISNYLNWYQQKPDGTVKLLIYYTSTLHSGVPSRFSGSGSGTDYSLTISNLEQEDVATYFCQ

**Linker** **17-4 VH**

QGYTLPWTFGGGTKLEIKGGGGSGGGGSGGGGSEVQLQQSGAELVKPGASVKLSCTASGFNIKETYIHWVKQRPEQGLEWIGRID

**Linker**

PADVYGRYDPKFQGKATITADTSSNSAYLQVSSLTSEDTAVYYCARSYGSSYGIDYWGQGTSVTVSSGGGGSLQQSGGGLVKPGGS

**IV.3 VH**

LKLSCAASGFTFSGYVMSWVRQSPEKRLEWVAEISSGGNYTYYPDTVTGRFTISRDNAKNTLYLEMNSLRSEDTAMYYCARVAYYG

**Linker** **IV.3 VL**

NYDYAMDYWGQGTSVTVSSGGGGSGGGGSGGGGSDIVLTQTTSSLSASLGDRVTISCRASQDITNYLNWYQQKPDGTLKLLIYYTS

**His-Tag**

RLHSGVPSRFSGSGSGTDYSLTISNLEQEDIATYFCQQGNTLRTFGGGTKLEIKRSRHHHHHH

#### (D) G6B monoclonal antibody 17.4 single-chain variable fragment

**17-4 VL**

DIQMTQTTSSLSASLGDRVTISCRASQDISNYLNWYQQKPDGTVKLLIYYTSTLHSGVPSRFSGSGSGTDYSLTISNLEQEDVATYFCQ

**Linker** **17-4 VH**

QGYTLPWTFGGGTKLEIKGGGGSGGGGSGGGGSEVQLQQSGAELVKPGASVKLSCTASGFNIKETYIHWVKQRPEQGLEWIGRID

**His-Tag**

PADVYGRYDPKFQGKATITADTSSNSAYLQVSSLTSEDTAVYYCARSYGSSYGIDYWGQGTSVTVSSHHHHHH

#### (E) GPVI monoclonal antibody 3J24 single-chain variable fragment

**3J24 VH**

EVQLQQSGGGLVKPGGSLKLSCAASGFTFSGYVMSWVRQSPEKRLEWVAEISSGGNYTYYPDTVTGRFTISRDNAKNTLYLEMNSL

**Linker** **3J24 VL**

RSEDTAMYYCARVAYYGNYDYAMDYWGQGTSVTVSSGGGGSGGGGSGGGGSDIVLTQTTSSLSASLGDRVTISCRASQDITNYLN

**His-Tag**

WYQQKPDGTLKLLIYYTSRLHSGVPSRFSGSGSGTDYSLTISNLEQEDIATYFCQQGNTLRTFGGGTKLEIKRSRHHHHHH

#### (F) CD32A monoclonal antibody IV.3 single-chain variable fragment

**IV.3 VH**

EIQLQQSGPELKKPGETVKISCKASGYTFTNYGMNWVKQAPGKGLKWMGWLNTYTGESIYPDDFKGRFAFSSETSASTAYLQINNL

**Linker** **IV.3 VL**

KNEDMATYFCARGDYGYDDPLDYWGQGTSVTVSSGGGGSGGGGSGGGGSDVVMTQTPPSVPVTPGESVSISCRSSKSLLHTNGN

**His-Tag**

TYLHWFLQRPGQSPQLLIYRMSVLASGVPDRFSGSGSGTAFTLSISRVEAEDVGVFYCMQHLEYPLTFGAGTKLELKHHHHHH

**Supplemental Figure 1: (A-F)** Amino acid sequences of recombinant G6B, GPVI and CD32A bi- and mono-specific single-chain variable fragments, as indicated.

### Supplemental Figure 2

**(A)**

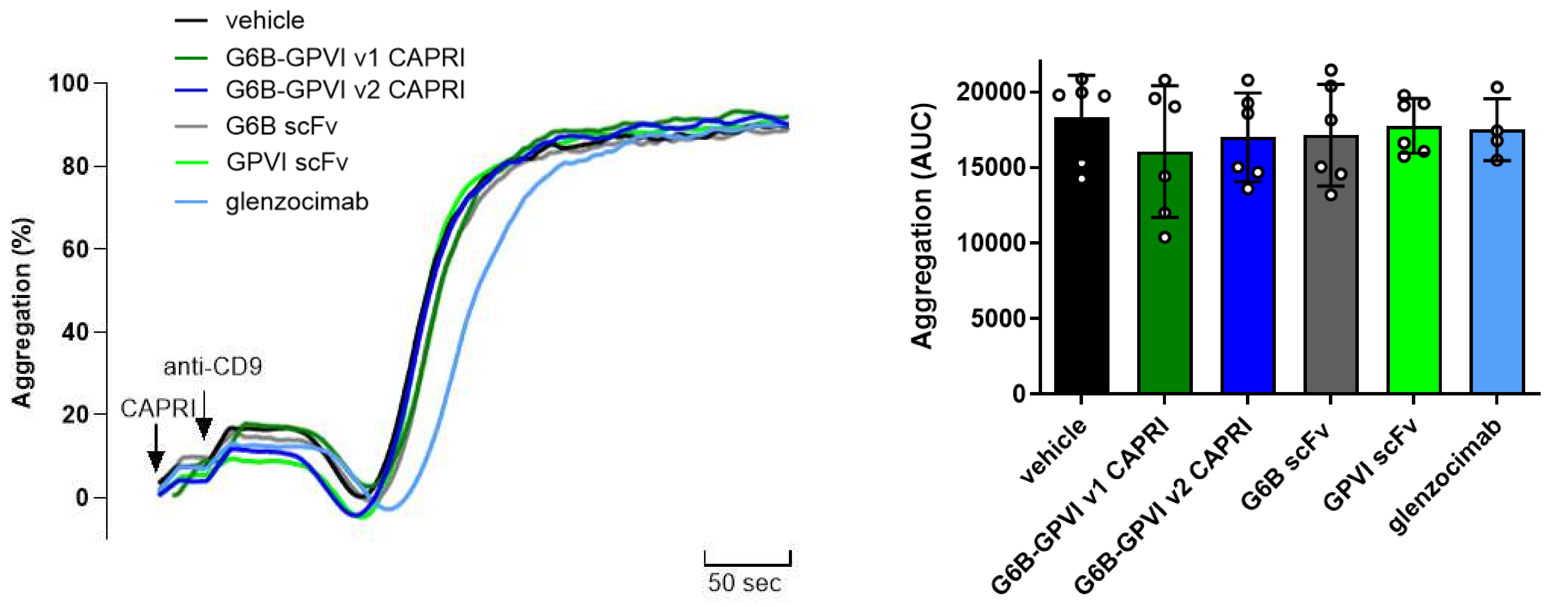


**(B)**

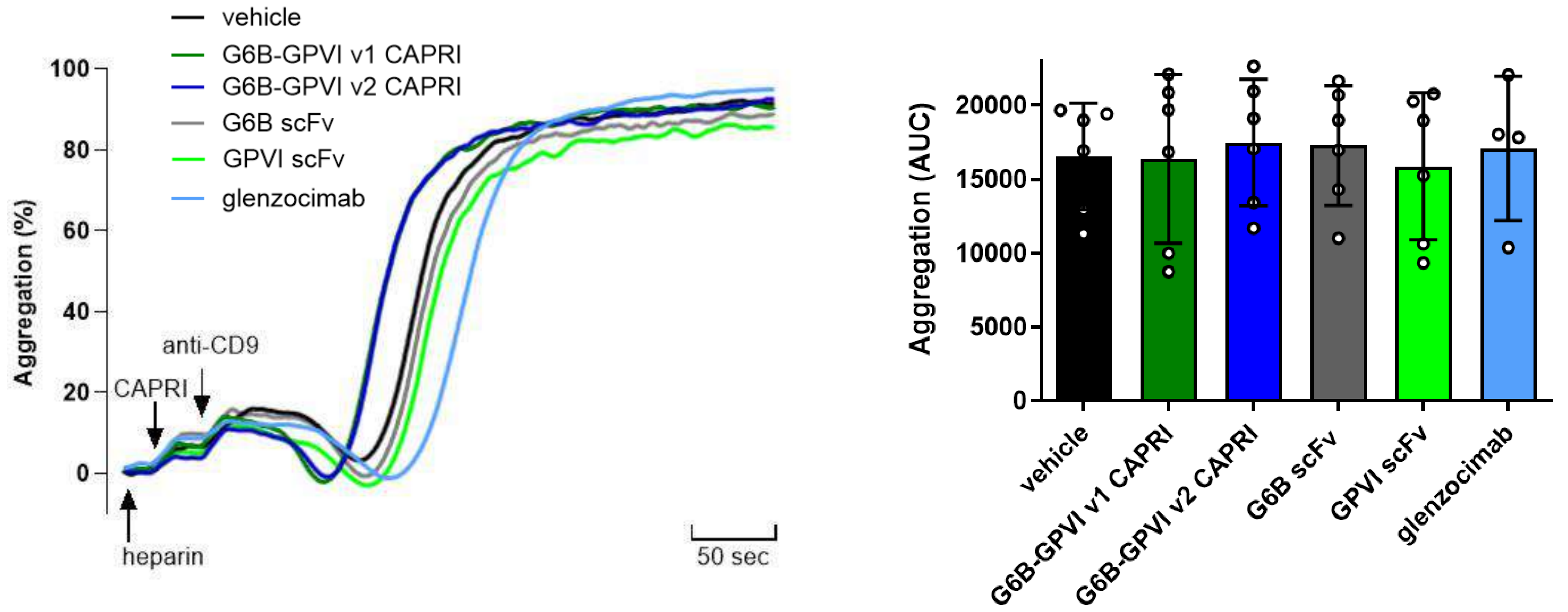


**Supplemental Figure 2. G6B-GPVI CAPRIs do not inhibit CD32A-mediated platelet aggregation.** Washed human platelets (3 x $10^8$/ml) from healthy donors, pre-treated with 0.2 μM of either G6B-GPVI v1, G6B-GPVI v2 CAPRIs, glenzocimab Fab, G6B scFv or GPVI scFv in the **(A)** absence and **(B)** presence of 1 U/ml heparin, prior to stimulation with 3 μg/ml of (20 nM) anti-CD9 mAb Alma1, which mediates platelet activation and aggregation in a CD32A-dependent manner. Platelet aggregation was measured in real-time using an APACT 4004 light transmission aggregometer, at 37°C with constant stirring. Representative traces and area under the aggregation curve (AUC) at 5 minutes are shown. (n = 4 per condition, mean ± SD).

### Supplemental Figure 3

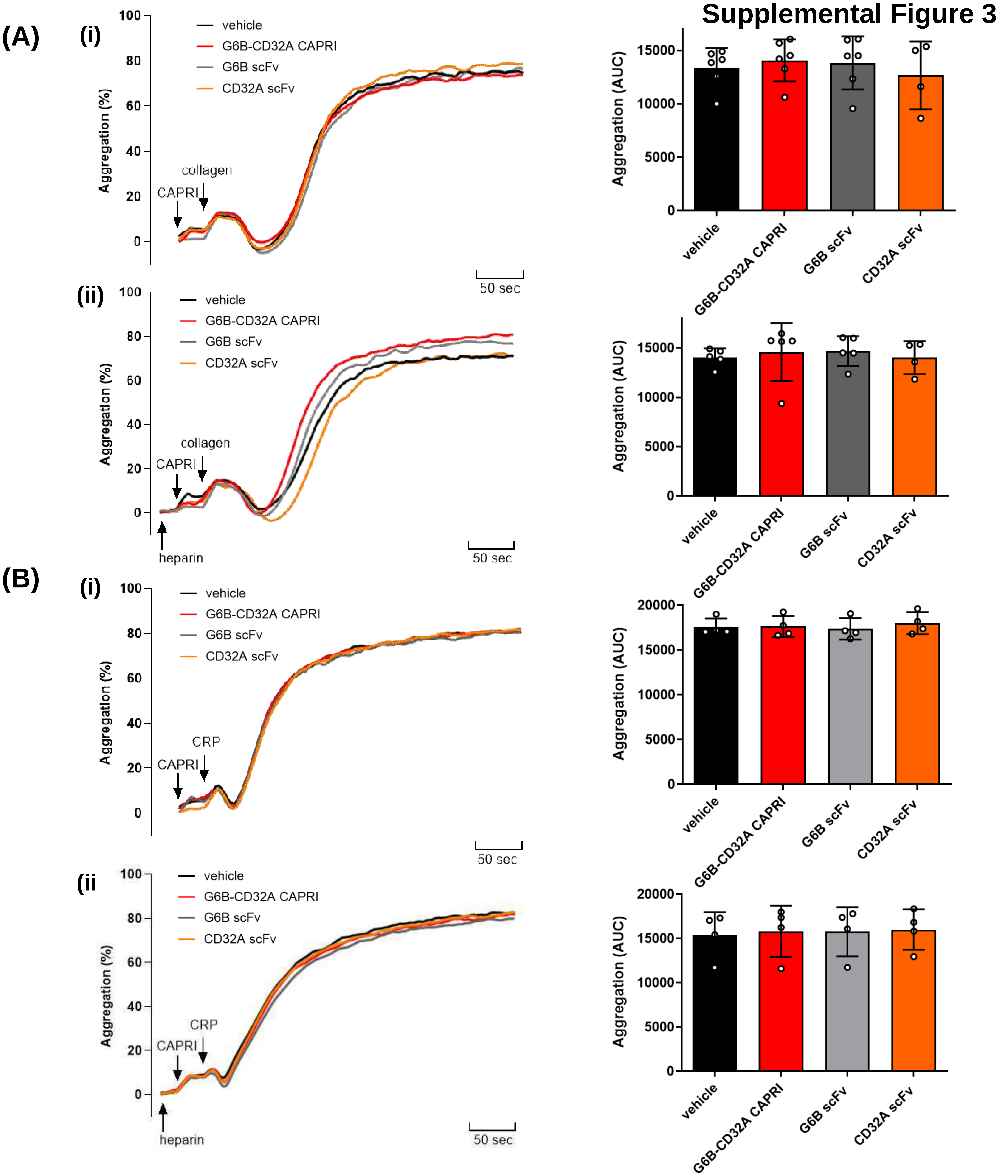


**Supplemental Figure 3. G6B-CD32A CAPRI does not inhibit collagen- and CRP-mediated platelet aggregation. (A)** Washed human platelets (3 x $10^8$/ml) from healthy donors were treated with either 0.2 μM G6B-CD32A CAPRI, G6B scFv or CD32A scFv, in the **(i)** absence and **(ii)** presence of 1 U/ml heparin, for 2 minutes at 37°C with constant stirring. Platelets were subsequently stimulated with **(A)** 3 μg/ml collagen or **(B)** 10 μg/ml CRP. Platelet aggregation was measured in real-time using an APACT 4004 light transmission aggregometer, at 37°C with constant stirring. Representative traces and area under the aggregation curve (AUC) at 5 minutes are shown. (n = 4-6 per condition, mean ± SD).

### Supplemental Figure 4

**(A)**

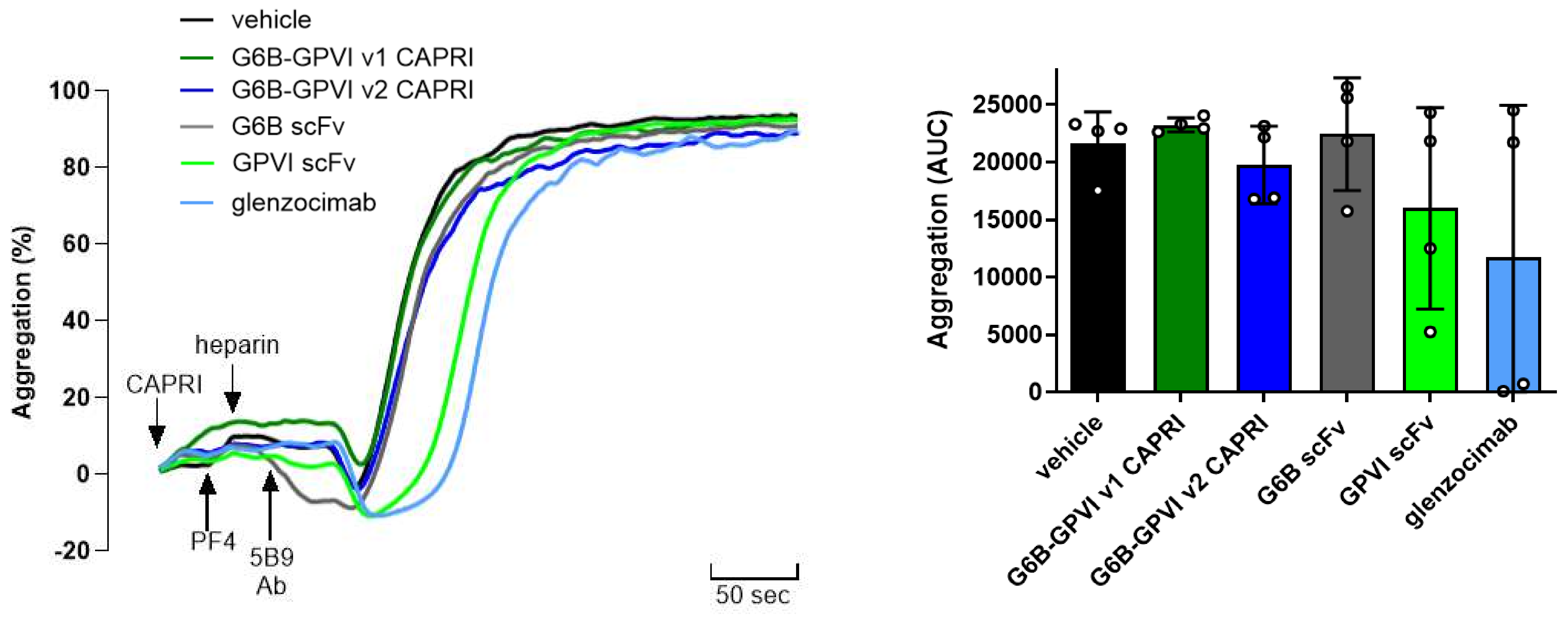


**(B)**

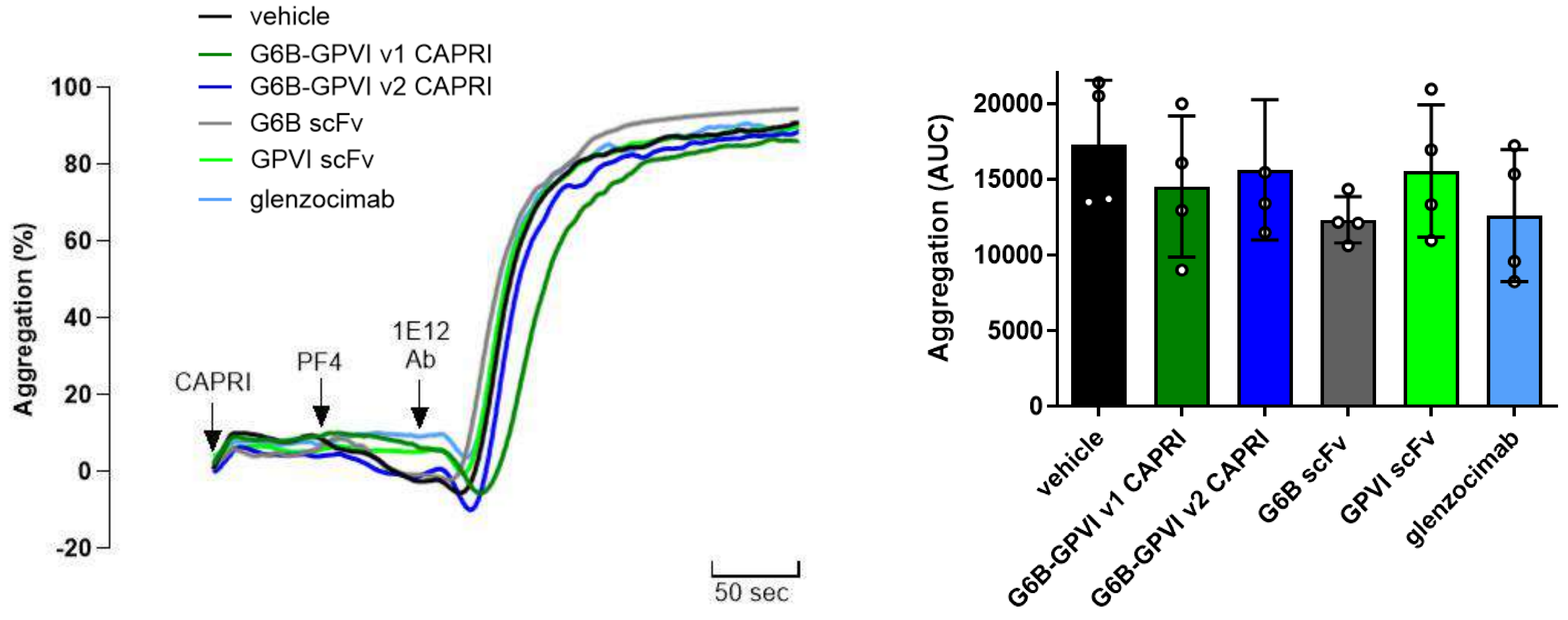


**Supplemental Figure 4. G6B-GPVI CAPRIs does not inhibit HIT- or VITT-mediated platelet aggregation. (A)** Washed human platelets (3 x $10^8$/ml) from healthy donors were treated with 1 mg/ml fibrinogen and either 0.2 µM G6B-GPVI v1, G6B-GPVI v2 CAPRIs, G6B scFv, GPVI scFv or glenzocimab, for 30 seconds at 37°C with constant stirring prior stimulation . Platelets were subsequently stimulated with a HIT immune complex consisting of PF4/heparin/mAb 5B9 by sequentially adding 10 µg/ml PF4, 0.5 U/ml heparin and 50 µg/ml 5B9 at 30-second intervals to platelet suspensions. **(B)** Same as in (A), but platelets were stimulated with a VITT rather than a HIT immune complex consisting of PF4/mAb 1E12 by sequentially adding 5 µg/ml PF4 and 10 µg/ml 1E12 at 1-minute intervals to platelet suspensions. Platelet aggregation was measured in real-time using an APACT 4004 light transmission aggregometer, at 37°C with constant stirring. Representative traces and area under the aggregation curve (AUC) at 5 minutes are shown. (n = 4 per condition, mean ± SD).

#### Supplemental Figure 5

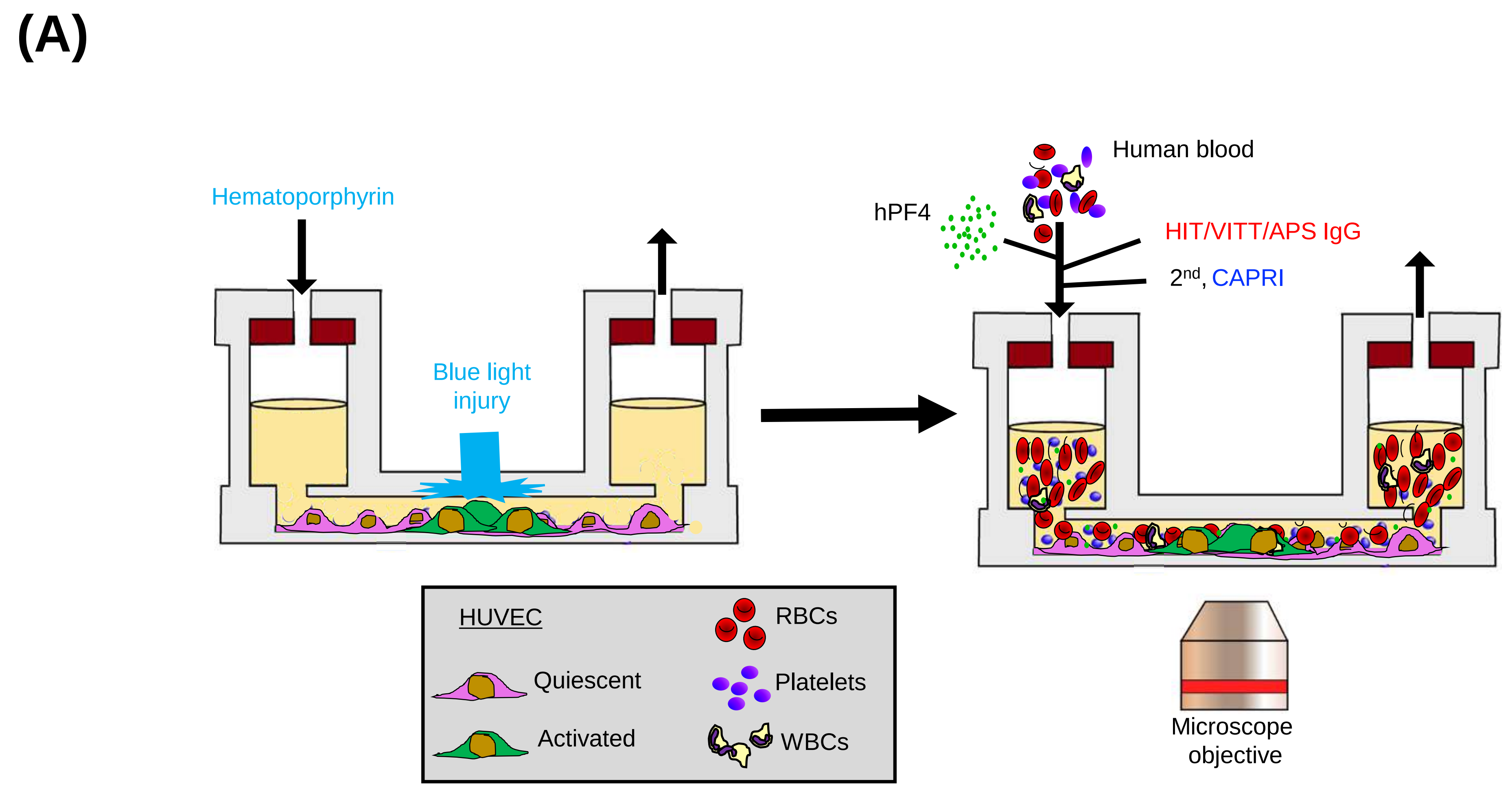


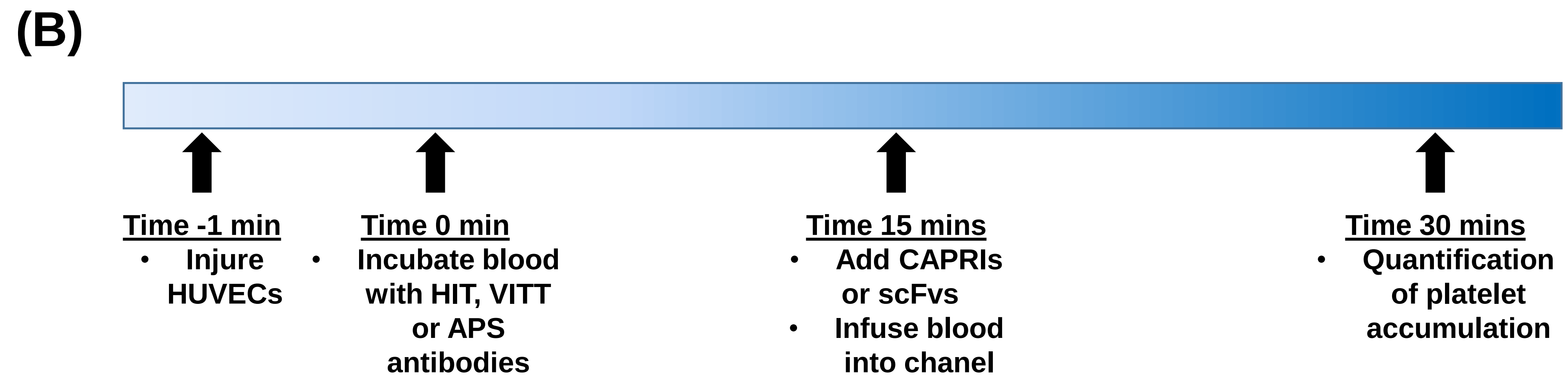


**Supplemental Figure 5. Schematics of the *in vitro* photochemical endothelial injury microfluidic HIT/VITT/APS model. (A)** shows the fibronectin-coated, HUVEC-lined system with whole blood flowed through status-post a hematoporphyrin-photochemical injury to part of the channel. Labeled-platelet accumulation during the study is the primary endpoint. **(B)** is the timeline for studies of infused products into the microfluidic channel experiments.

### Supplemental Figure 6

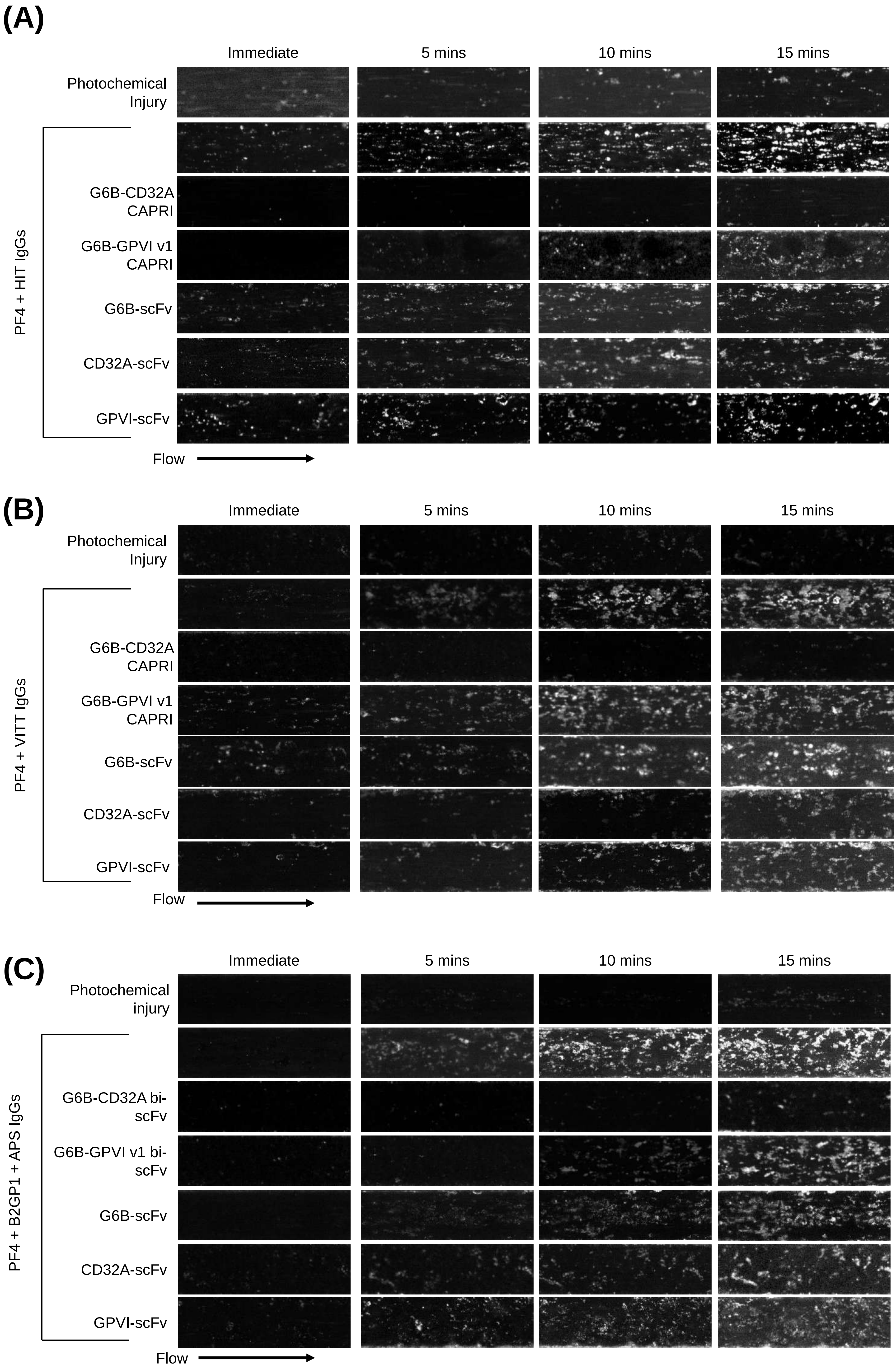


**Supplemental Figure 6. Effect of CAPRIs on HIT/VITT/APS thrombosis in a microfluidic system.** Representative images over a 15-minute window of flow through the channels showing platelet accumulation (white aggregates) on the endothelial lining, following **(A)** HIT, **(B)** VITT or **(C)** APS antibody-induced activation.
