## Supplemental Material and Methods for "G6b-B antibody-based *cis*-acting platelet receptor inhibitors (CAPRIs) as a new family of anti-thrombotic therapeutics"

### Supplemental Materials and Methods

#### Antibodies and reagents

Anti-Shp1, anti-Src, anti-Syk, anti-Shp1 *p*-Tyr564, anti-Shp2 *p*-Tyr542 and anti-Syk *p*-Tyr525/526 antibodies were from Cell Signaling Technology. Anti-Src *p*-Tyr418 antibody was from Thermo Fisher Scientific and anti-Shp2 antibody from Santa Cruz **(Supplemental Table 2)**. Collagen-related peptide (CRP)-XL was obtained from Triple Helical Peptides (Cambridge, UK). Collagen was from Takeda (Horm, Linz, Austria). Heparin Choay was obtained from Sanofi, FR.

#### Production of bi-scFv CAPRIs

Bi- and mono-specific scFvs were generated by Peak Proteins (Sygnature Discovery, Macclesfield, UK). The light- and heavy-chain variable sequences (VL and VH, respectively) of anti-G6B mAb 17-4 were recombinantly joined to the VL and VH of either anti-GPVI mAbs 3J24 or glenzocimab (Fab of humanized anti-GPVI mAb ACT017),(1) or anti-CD32A mAb IV.3,(2) using flexible glycine (G)/serine (S)-rich linker peptides, resulting in three distinct bi-scFv CAPRIs: (i) G6B-GPVI version 1 (v1), (ii) G6B-GPVI v2, and (iii) G6B-CD32A **(Figures 1A and 1B, Supplemental Figures 1A-F and Table 1)**. Nucleotide sequences were codon optimized for expression in mammalian cells. The N-termini of each (bi-)scFv was engineered with an Ig kappa leader sequence (METDTLLLWVLLLWVPGSTG) to enable secretion from mammalian cells, and C-terminal 6-histidine (His-)tag to facilitate protein purification. scFvs were used as negative controls.

Synthetic DNAs of (bi-)scFvs were sub-cloned into the pTT5 mammalian expression vector and transiently expressed in HEK293 cells at 2 and 4 liter scale. His-tagged proteins were purified using nickel affinity and size exclusion chromatography. Protein purity was determined by sodium dodecyl sulphate-polyacrylamide gel electrophoresis (SDS-PAGE) and size-exclusion chromatography, and identities confirmed by mass spectrometry. Protein concentrations were determined by absorbance at 280 nm. Expected and observed molecular masses of bi-scFvs and scFvs were approximately 54 and 27 kDa, respectively.

**Platelet preparation and aggregation**

Biological activities of G6B-GPVI and G6B-CD32A CAPRIs and corresponding scFv and Fab controls were tested at equimolar concentrations by standard light transmission platelet aggregation, using an APACT 4004 platelet aggregometer. All steps were performed at 37°C with constant stirring at 1,200 rpm. Briefly, blood was collected from healthy donors into anticoagulant citrate-dextrose solution (ACD), as previously described.(3) Washed human platelets were prepared using a standard protocol and suspended in modified Tyrode's buffer at a concentration of $3 \times 10^8$/ml, as previously described.(3) Platelets were pre-treated with either 10 μg/ml (0.2 μM) of G6B-GPVI v1, v2, or G6B-CD32A CAPRIs, CD32A Fab, glenzocimab, G6B, CD32A, or GPVI scFvs, prior to stimulation with the either 10 μg/ml of the GPVI-specific agonist collagen-related peptide (CRP), 3 μg/ml collagen or 3 μg/ml (20 nM) of the anti-CD9 mAb Alma1, which mediates platelet activation via CD32A.(4)

Platelets were also stimulated with HIT and VITT immune complexes. In the case of HIT, either 100 μg/ml (645 nM) KKO mAb (R&D Systems), or generated as described,(5) or 50 μg/ml (323 nM) 5B9 mAb was added to washed platelets in suspension at 37°C with constant stirring, after 1 mg/ml (2.4 mM) fibrinogen, 10 μg/ml recombinant human PF4 (R&D Systems), or generated as described,(5, 6) and 0.5 U/ml heparin. VITT immune complexes were formed by first adding 1 mg/ml (2.4 mM) fibrinogen to washed platelets in suspension at 37°C with constant stirring, followed by 5 μg/ml (43 nM) recombinant human PF4, and 10 μg/ml (65 nM) 1E12 mAb, generated as described.(7)

**Platelet stimulation for biochemical analysis**

For biochemical analysis of GPVI and CD32A signaling pathways, platelets were pre-incubated with ReoPro (0.2 μM), apyrase (2 Unit/ml) and indomethacin (10 μM), and with either 10 μg/ml (0.2 μM) of G6B-GPVI v1, v2, or G6B-CD32A CAPRIs, CD32A Fab, glenzocimab, G6B, CD32A, or GPVI scFvs for 10 minutes, under stirring conditions (1,200 rpm, 37°C), prior stimulation with CRP or anti-CD9 mAb Alma1 for 90 seconds.

**Immunoblotting**

Platelet whole cell lysates (WCLs) were prepared and analyzed by automated capillary-based immunoassay (ProteinSimple Jess), prepared according to manufacturer's instructions and as previously described.(8) Briefly, WCLs were diluted to the required concentration with 0.1× sample buffer, followed by addition of 5× master mix containing 200 mM dithiothreitol (DTT), 5× sample buffer, fluorescent standards and boiled for 5 minutes at 95°C. Samples containing antibody diluent, primary and anti-rabbit secondary antibodies, luminol S-peroxide mix, stripping buffer and wash buffer, were loaded on Jess 12-230 kDa prefilled microplates. Primary antibodies were incubated and the High Dynamic Range (HDR) profile was used for chemiluminescent and fluorescent multiplexing signal detection. Optimized antibody dilutions and sample concentrations used are provided in **Supplemental Table 2**.

**Collagen microfluidic thrombosis studies**

Platelet adhesion and thrombus formation was measured using a microfluidic system, as previously described.(9) Briefly, polydimethylsiloxane (PDMS) flow chambers channels (0.1 × 1mm) were coated with collagen (200 µg/ml) for 1 hour at room temperature and blocked with phosphate-buffered saline (PBS)-containing human serum albumin (10 mg/ml) for 30 minutes at room temperature. Whole blood was collected from healthy donors into 525 U/ml hirudin and treated with 0.2 µM of the various (bi-)scFvs for 15 minutes at 37°C prior to being perfused through the collagen-coated chamber with a syringe pump (Harvard Apparatus, Holliston, MA, USA) at 1,500 $s^{-1}$ for 5 minutes at 37°C. Platelets adhesion and aggregation were visualized in real-time by differential interference contrast (DIC) microscopy (Leica DMI4000B). Platelets were also stained with 3,3′-dihexyloxacarbocyanine iodide ($DiOC_6$) and visualized by confocal microscopy. Surface coverage by platelet aggregates (>100 $µm^2$) and thrombus volume were quantified using ImageJ software. All experiments were performed blind.

**HIT and VITT microfluidic thrombosis studies on VWF**

Microfluidic thrombosis models were performed as previously described.(7) Briefly, whole blood was collected on 0.129 M sodium citrate from healthy donors. It was diluted in half with PBS and pre-incubated with 0.1 µM of the various (bi-)scFvs for 10 minutes, prior to incubation with 1E12 at 10 µg/mL for 6 minutes. Blood samples were then recalcified to 2.7 mM $CaCl_2$ and perfused at a shear rate of 20 µL/min (500 $s^{-1}$; 20 dyn/$cm^2$) in microfluidic channels (Vena8 Fluoro+, Cellix), pre-coated overnight at 4°C with 120 µg/mL purified human VWF (LFB). Leukocytes were visualized by adding Hoechst (16.2 µM). Cellular aggregates (leukocytes and platelets) and fibrin deposition were observed by staining with 17.4 µM $DiOC_6$ and 44 nM AF647-fibrinogen (Invitrogen), respectively. An Axio Observer 7 microscope (Zeiss) and a

LD Plan-Neofluar 20x/0.4 Ph2 objective equipped with an ORCA Flash 4.0 LT plus C11440 digital CCD camera (Hamamatsu) controlled by Zen 2.6 2018 image capture software were used to acquire pictures. Whole blood was perfused for 8 minutes and pictures were taken at the end of the perfusion. The final images are used to measure the area covered with aggregates (> 100 $\mu m^2$) using ImageJ software.

**Endothelial injury HIT, VITT and APS microfluidic studies**

Microfluidic studies in 48-well BioFlux plates (Fluxion biosciences) were performed with channels coated with near-confluent human umbilical vein endothelial cells (HUVECs, Lifeline Cell Technology), as previously described.(5) HUVECs were injured using an HXP120C light source with a 475-nm excitation and 530-nm emission filter for 20 seconds while the channel was perfused with 50 µg/ml hematoporphyrin (Sigma-Aldrich). Whole blood collected in 0.32% citrate was labeled with 2 mM calcein AM green (ThermoFisher) and supplemented with recombinant PF4 (25 µg/ml) and either 1 mg/ml of HIT, VITT or APS IgGs isolated from patient samples. After 15 minutes, 0.6 µM of the various (bi-)scFvs were added and samples perfused into channels at 10 dyne/$cm^2$. Images were captured 5, 10, and 15 minutes after perfusion was initiated. Platelet accumulation on the injured endothelium field was captured using a Zeiss Axio Observer Z1 inverted microscope using Montage Fluxion software and analyzed using ImageJ. Confluent, uninjured endothelial-lined channels exposed to blood with no added PF4 and no HIT, VITT and APS antibodies were used as the background, and was subtracted from values obtained from injured vessels exposed to blood that had been exposed to various experimental conditions.

### Statistical analysis

The set of variables is described using the usual position and dispersion parameters, namely mean, median, quartiles, minimum and maximum, as well as standard deviation and variance. The Gaussian character of a quantitative variable was assessed using the Shapiro-Wilk normality test. All data is presented as mean ± standard deviation (SD). Statistical significance was analyzed by one- or two-way ANOVA followed by the appropriate *post hoc* test, or Kruskal-Wallis test for nonparametric data, a Bonferroni correction was applied for multiple comparisons, as indicated in figure legends. All analyses were performed using GraphPad Prism9 software (GraphPad Software Inc, San Diego, CA, US). Differences were considered significant when the P values were $<0.05$.

**Supplemental Table 1: Bispecific single-chain variable fragments and controls**

| Reagents | Abbreviations |
|---|---|
| Anti-G6B mAb 17.4 scFv | G6b scFv |
| Anti-GPVI mAb 3J24 scFv | GPVI scFv |
| Anti-CD32A mAb IV.3 scFv | CD32A scFv |
| Anti-CD32A mAb IV.3 Fab | CD32A Fab |
| Anti-GPVI mAb ACT017 Fab | glenzocimab |
| Anti-G6B mAb 17.4 VR-GPVI mAb 3J24 VR bi-scFv | G6B-GPVI v1 CAPRI |
| Anti-G6B mAb 17.4 VR-GPVI mAb ACT017 VR bi-scFv | G6B-GPVI v2 CAPRI |
| Anti-G6B mAb 17.4 VR-CD32A mAb IV.3 VR bi-scFv | G6B-CD32A CAPRI |

Single-chain variable fragments (scFv)
Fragment antigen-binding (Fab)
Variable region (VR)
Bispecific (bi-)

**Supplemental Table 2: Antibody dilutions and sample concentrations used in ProteinSimple Jess**

| Antibody | Reference | Dilution | Sample (mg/ml) |
|---|---|---|---|
| Rabbit anti-Syk *p*-Tyr525/526 | 2711 CST | 1/10 | 0.2 |
| Rabbit anti-Syk | 12358 CST | 1/50 | 0.2 |
| Rabbit anti-Src *p*-Tyr418 | 44-660 TFS | 1/25 | 0.1 |
| Rabbit anti-Src | 2108 CST | 1/25 | 0.1 |
| Rabbit anti-Shp2 *p*-Tyr542 | 3703 CST | 1/25 | 0.2 |
| Mouse anti-Shp2 | SC-7384 SC | 1/10 | 0.2 |
| Rabbit anti-Shp1 *p*-Tyr564 | 8849 CST | 1/25 | 0.1 |
| Rabbit anti-Shp1 | 3759 CST | 1/10 | 0.1 |

Cell Signaling Technology (CST)
Thermo Fisher Scientific (TFS)
Santa Cruz (SC)

**Supplemental Video 1:** Whole, hirudinated blood collected from healthy donors was flowed over a collagen-coated at 1,500 $s^{-1}$ for 5 minutes at 37°C. The accumulation of adherent platelets on collagen fibrils was monitored in real-time using DIC microscope. The time in seconds is indicated. Representative video from 5 independent experiments.

**Supplemental Video 2:** Whole, hirudinated blood collected from healthy donors was treated with 0.2 μM of G6B-GPVI v1 CAPRI for 15 minutes at 37°C prior to being flowed over a collagen-coated at 1,500 $s^{-1}$ for 5 minutes at 37°C. The accumulation of adherent platelets on collagen fibrils was monitored in real-time using DIC microscope. The time in seconds is indicated. Representative video from 5 independent experiments.

**Supplemental Video 3:** Whole, hirudinated blood collected from healthy donors was treated with 0.2 μM of G6B-GPVI v2 CAPRI for 15 minutes at 37°C prior to being flowed over a collagen-coated at 1,500 $s^{-1}$ for 5 minutes at 37°C. The accumulation of adherent platelets on collagen fibrils was monitored in real-time using DIC microscope. The time in seconds is indicated. Representative video from 5 independent experiments.

**Supplemental Video 4:** Whole, hirudinated blood collected from healthy donors was treated with 0.2 μM of G6B-CD32A CAPRI for 15 minutes at 37°C prior to being flowed over a collagen-coated at 1,500 $s^{-1}$ for 5 minutes at 37°C. The accumulation of adherent platelets on collagen fibrils was monitored in real-time using DIC microscope. The time in seconds is indicated. Representative video from 5 independent experiments.

**Supplemental Video 5:** Whole, hirudinated blood collected from healthy donors was treated with 0.2 μM of G6B scFv for 15 minutes at 37°C prior to being flowed over a collagen-coated at 1,500 $s^{-1}$ for 5 minutes at 37°C. The accumulation of adherent platelets on collagen fibrils was monitored in real-time using DIC microscope. The time in seconds is indicated. Representative video from 5 independent experiments.

**Supplemental Video 6:** Whole, hirudinated blood collected from healthy donors was treated with 0.2 μM of GPVI scFv for 15 minutes at 37°C prior to being flowed over a collagen-coated at 1,500 $s^{-1}$ for 5 minutes at 37°C. The accumulation of adherent platelets on collagen fibrils was monitored in real-time using DIC microscope. The time in seconds is indicated. Representative video from 5 independent experiments.

**Supplemental Video 7:** Whole, hirudinated blood collected from healthy donors was treated with 0.2 μM of glenzocimab scFv for 15 minutes at 37°C prior to being flowed over a collagen-coated at 1,500 $s^{-1}$ for 5 minutes at 37°C. The accumulation of adherent platelets on collagen fibrils was monitored in real-time using DIC microscope. The time in seconds is indicated. Representative video from 5 independent experiments.

**Supplemental Video 8:** Whole, hirudinated blood collected from healthy donors was treated with 0.2 μM of CD32A scFv for 15 minutes at 37°C prior to being flowed over a collagen-coated at 1,500 $s^{-1}$ for 5 minutes at 37°C. The accumulation of adherent platelets on collagen fibrils was monitored in real-time using DIC microscope. The time in seconds is indicated. Representative video from 5 independent experiments.
